# Region-specific molecular regulatory programs define epithelial identity, progenitor states, and mucus homeostasis in human distal airways

**DOI:** 10.64898/2026.06.15.732400

**Authors:** Hiroaki Murano, Nathanial C. Stevens, Hong Dang, Satoko Nakano, Anne M. Cawley, Zoey Y. Wisniewski, Gillian Crisp, Elodie Mitchell, Avishi Singh, Sindhuja Damodaran, Boris Reidel, Mark I. Gutay, Ling Sun, Koichi Hasegawa, Allison B. Williams, Robert M. Immormino, Rodney C. Gilmore, Lisa C. Morton, Minako Furusho, Takanori Asakura, Yu Mikami, Thomas Whitlow, Naoya Miyashita, Rhianna E. Lee-Ferris, Nancy L. Quinney, Susan E. Boyles, Deborah M. Cholon, Michael Chua, Takafumi Kato, Leslie Fulcher, Hirotoshi Matsui, Gang Chen, Alessandra Livraghi-Butrico, Martina Gentzsch, Stephen A. Schworer, Brian Button, Mehmet Kesimer, Purushothama Rao Tata, James S. Hagood, Gloria S. Pryhuber, Wanda K. O’Neal, Scott H. Randell, Richard C. Boucher, Kenichi Okuda

## Abstract

Small distal airways differ from proximal large airways in structure, airflow dynamics, and epithelial composition, and represent a central site of muco-obstructive lung disease pathogenesis. However, due in part to their inaccessibility, the molecular mechanisms that establish regional epithelial identity and govern mucociliary defense in distal airway epithelia remain poorly defined. Here, we integrate transcriptomic, secretomic, and chromatin accessibility analyses of matched primary human large and small airway epithelial cultures to define region-specific regulatory networks. We identify distal airway-specific transcriptional and chromatin programs required for maintaining epithelial identity and mucus homeostasis. Loss of NKX2-1 impairs distal airway secretory cell (DASC) differentiation and shifts mucus properties toward a disease-associated state. Lineage-resolved organoid assays identify an NKX2-1-high distal airway basal cell population with hybrid basal-secretory features as a selective progenitor for DASCs. Collectively, these findings establish a molecular framework for distal airway epithelial biology and define mechanisms regulating region-specific mucociliary host defense.

## Introduction

Most human muco-obstructive lung diseases (MOLDs), including asthma^1, 2^, cystic fibrosis (CF)^3–9^, and chronic obstructive pulmonary disease (COPD)^10, 11^, originate in and are dominated by “small” airways, i.e., the bronchioles^1, 3–9^. Traditionally defined as airways less than 2 mm in diameter and lacking submucosal glands and cartilage, bronchioles differ from proximal tracheobronchial airways in anatomical structures^12^, airflow dynamics^13^, and epithelial cell compositions^14–19^. Owing to the highly branched architecture of the airway tree, bronchioles vastly outnumber large airways and account for the majority of conducting airway surface area^20^, highlighting their central role in lung function and innate lung defense. Despite their significance, the molecular mechanisms by which bronchiolar epithelial cells establish and maintain their distinct regional identity, and, importantly, how this identity supports distal airway mucosal homeostasis, remain poorly defined. In part, this gap reflects the absence of physiologically relevant model systems that enable mechanistic and functional interrogation of distal airway epithelial biology.

Recent spatial and single-cell transcriptional profiling studies have revealed striking epithelial cell diversity along the proximal-distal axis of the human lung^14–19^. A distal bronchiole-specific secretory cell population, variably termed respiratory airway secretory (RAS) cells^17^, terminal airway-enriched secretory cells (TASCs)^18^, or pre-terminal/terminal respiratory bronchiolar secretory cells (pre-TB/TRB-SC)^19^, has been identified as a distinct regional cell type characterized by expression of *SFTPB* and *SCGB3A2*. This cell type, which we refer to as the distal airway secretory cell (DASC), exhibits multipotent progenitor properties, including transdifferentiation into both airway and alveolar epithelial cells^19^. Beyond their progenitor function, many aspects of DASC biology remain poorly understood, including their role in mucociliary clearance (MCC) and host defense, the transcriptional programs that maintain their region-specific identity, and their upstream lineage origins. Importantly, phenotypic loss of DASCs has been observed in multiple MOLDs, including asthma^21^, COPD^16, 18^, and idiopathic pulmonary fibrosis (IPF)^22^, all of which exhibit impaired distal airway MCC^18, 22–24^. These observations suggest that DASCs play a critical role in maintaining distal airway mucociliary host defense and that regulatory programs specifying DASC differentiation are disrupted in disease.

In this study, we leveraged matched primary large airway epithelial (LAE) and small airway epithelial (SAE) cultures from a total of 57 donor lungs to define region- and cell-type-specific epithelial transcriptional programs, secretory proteomes, chromatin regulatory landscapes, and progenitor functions. Integrative multiomic analyses uncovered NKX2-1-dependent, region-specific molecular regulatory programs that were required for DASC differentiation and mucus homeostasis. We further identified a previously unrecognized lineage relationship between a distal airway-specific basal cell population and DASCs. Collectively, these studies establish a molecular framework for human distal airway epithelial identity and provide a platform for multimodal investigation of distal airway epithelial biology and pathophysiology.

## Results

### Region-specific transcriptomic and secretory programs in human airway epithelia

To define intrinsic regional identity in human airway epithelia, we established matched primary LAE and SAE cultures from previously healthy donor lungs (**Supplementary Table 1-2**) using conditionally reprogrammed cell (CRC) expansion methods^15, 25^, followed by air-liquid interface (ALI) differentiation (**Fig. 1a**). SAE cultures selectively expressed canonical DASC markers (SFTPB, SCGB3A2) (**Fig. 1b**), consistent with their bronchiole-restricted localization in vivo (**Fig. 1c**), whereas these markers were absent from LAE. These markers, together with *SCGB1A1*, were consistently enriched in SAE across differentiation, while major epithelial lineage markers, including mucin-secreting (*MUC5B*, *MUC5AC*), ciliated (*FOXJ1*), basal (*TP63*), and ionocytes (*FOXI1*), were comparably expressed between regions (**Fig. 1d**).

**Fig. 1.**
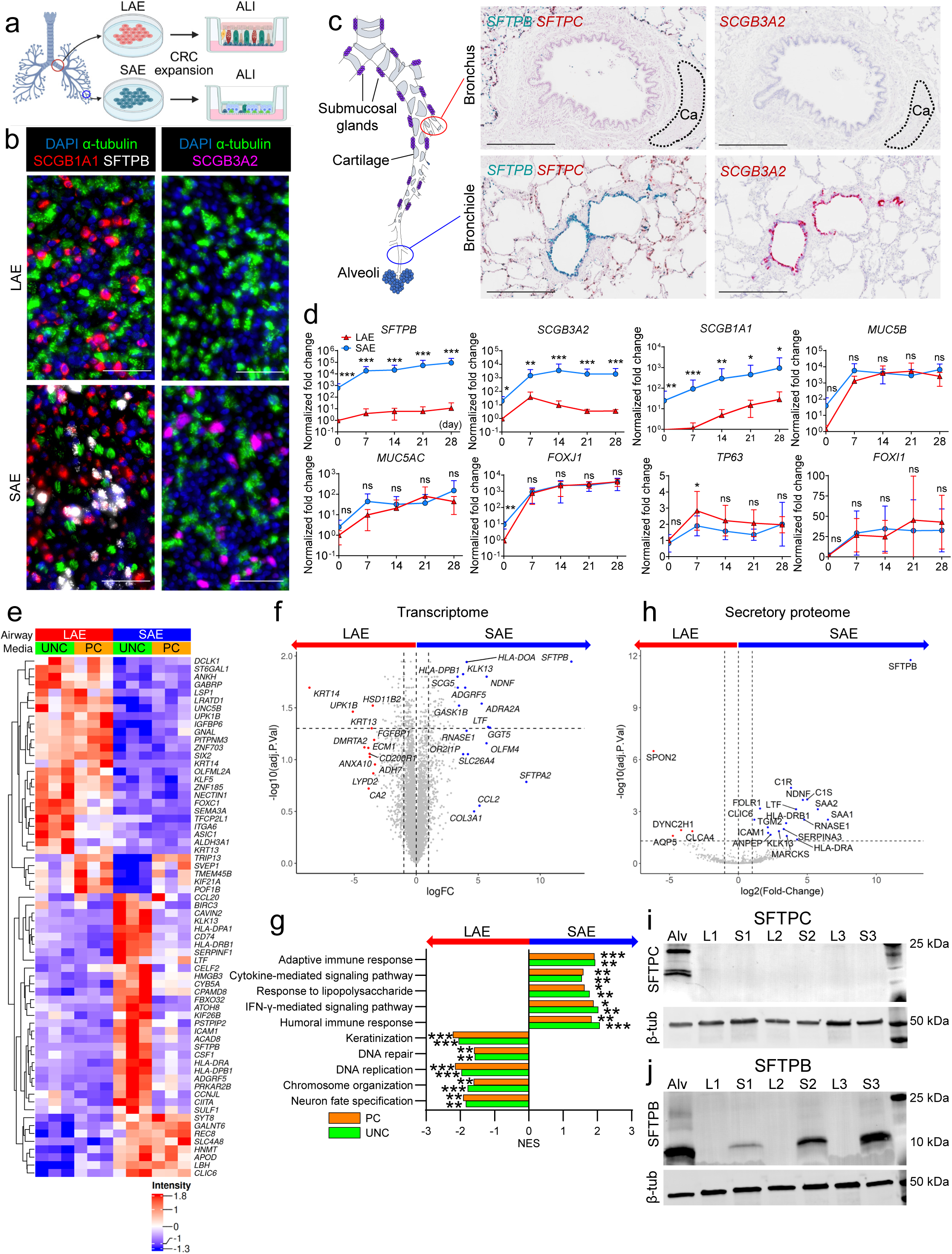
Transcriptomic and secretory proteomic characterization of primary human LAE and SAE. **a.** Schematic of LAE and SAE air-liquid interface (ALI) cultures. **b.** Immunofluorescence for epithelial markers in LAE and SAE ALI cultures. Scale bars, 50 μm. **c.** RNA *in situ* hybridization in a normal human lung. Ca, cartilage. Scale bar, 500 μm. **d.** Transcript expression during ALI differentiation (n = 5∼6). **e.** Differentially expressed genes (DEGs) between LAE and SAE across two distinct culture media. UNC, UNC ALI medium; PC, PneumaCult ALI medium. **f.** Top 30 DEGs ranked by fold change between LAE and SAE across combined media conditions. **g.** Gene set enrichment analysis comparing LAE and SAE across combined media conditions. **h.** Differentially enriched secretory proteins in LAE and SAE secretions (combined microdissection and bulk digestion datasets). **i, j.** Western blot of SFTPC (**i**) and SFTPB (**j**) in LAE (L), SAE (S), and alveolar epithelial (Alv) cultures. n = 3. ns, not significant; *p < 0.05; **p < 0.01; ***p < 0.001 (linear mixed-effects model).

Bulk RNA-seq demonstrated that regional identity was preserved across culture conditions, including distinct media (UNC ALI and PneumaCult™ ALI media), although culture media influenced epithelial morphology and select transcriptional programs (**Fig. 1e, Extended Data Fig. 1a-e**). Epithelial height in LAE remained greater than in SAE across media conditions, recapitulating in vivo morphologies (**Extended Data Fig. 1b**). LAE cultures were enriched for epithelial barrier and keratinization programs *(KRT14*, *KRT13*, *UPK1B*), whereas SAE cultures preferentially expressed DASC-associated and host defense genes (*SFTPB*, *SCGB3A2*, *APOD*, *LTF*, *KLK13*, *RNASE1*) (**Fig. 1f-g, Extended Data Fig. 1f**).

Consistent with these transcriptional differences, mass spectrometry of apical secretions revealed a distinct and more diverse secretory proteome in SAE, including SFTPB, RNASE1, KLK13, and LTF (**Fig. 1h**), while SCGB3A2 was not detected under these conditions. These distal airway-enriched secretory features were preserved across SAE cultures derived using two independent approaches to isolate distal airway progenitor cells, including microdissection and bulk enzymatic digestion (**Extended Data Fig. 2**), indicating that region-specific epithelial lineage differentiation is intrinsically encoded within distal airway progenitor cells, independent of isolation methods.

Supporting the specificity of the SAE cultures, pro-SFTPC was undetectable in both LAE and SAE lysates but remained robustly expressed in cultured alveolar epithelial type 2 (AT2) cells used as positive control^26, 27^, indicating minimal AT2 contamination in SAE despite their anatomical proximity (**Fig. 1i**). In contrast, SFTPB was readily detected in SAE and AT2 lysates, but not in donor-matched LAE (**Fig. 1j**). Together, these findings establish that SAE retain a molecularly and functionally distinct identity from both proximal airway and alveolar epithelial cells.

### Single-cell mapping of regional airway epithelial heterogeneity

Bulk RNA-seq and secretory proteomics revealed robust region-specific transcriptional and secretory signatures in LAE versus SAE cultures. Notably, SAE secretions contained a larger set of regionally enriched proteins than LAE secretions (**Fig. 1h, Extended Data Fig. 2e**), highlighting the specialized secretory function of bronchiolar epithelial cells. To resolve cell type-specific contributions to these regional differences, we performed single cell RNA sequencing (scRNA-seq) on differentiated LAE and SAE cultures from five matched non-disease donor lungs (**Fig. 2a, Extended Data Fig. 3a-c**). Uniform manifold approximation and projection (UMAP) analysis identified diverse epithelial cell clusters shared across LAE and SAE (**Fig. 2b-d, Extended Data Fig. 3d**). After low-quality clusters were excluded based on low numbers of detected genes and total transcript counts (**Extended Data Fig. 3e-g**), clusters were defined based on canonical marker gene expression, irrespective of regional origin (**Fig. 2e**). Differentially expressed gene (DEG) analyses of LAE and SAE within each cluster revealed distinct regionally enriched transcriptional signatures (**Extended Data Fig. 3h-m**). Notably, among SAE-derived cell populations, SAE-enriched genes identified by bulk RNA-seq (*SFTPB*, *SCGB3A2*, *RNASE1*, *ADGRF5*) (**Fig. 1e-f, Extended Data Fig. 1f**) were concentrated in secretory and hybrid secretory-ciliated cells, whereas basal, suprabasal, and cycling cells primarily expressed LAE-enriched genes (*KRT13*, *KRT14*, *UPK1B*) (**Fig. 2e-f**). These findings indicate that the strongest region-specific transcriptional divergence occurs in secretory and basal cell populations.

**Fig. 2.**
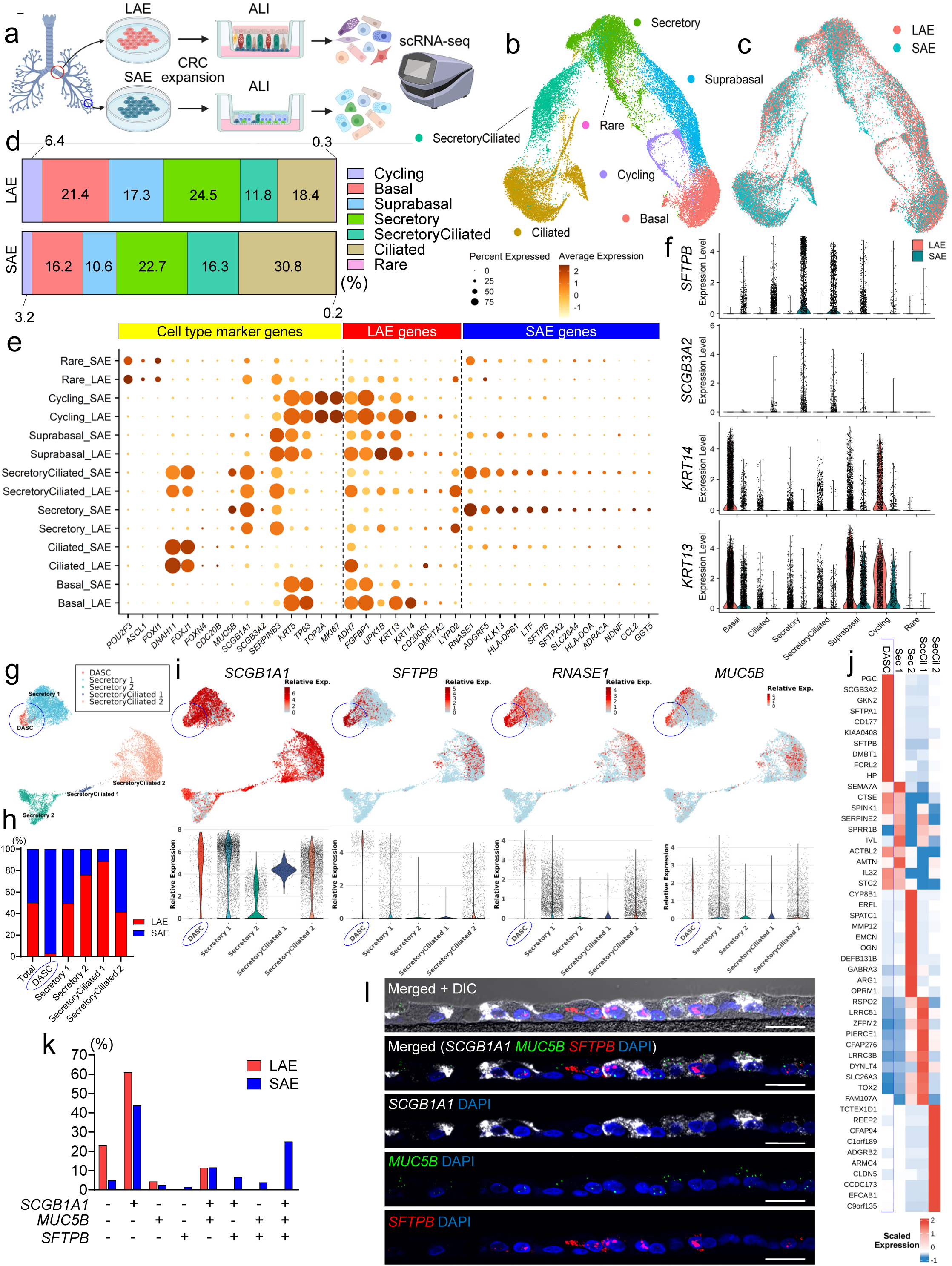
Identification of a distal airway-specific secretory cell population defining SAE identity. **a.** Experimental design. **b, c.** UMAP of epithelial cells from LAE and SAE (31,017 cells; n = 5) (**b**), colored by regional origin (**c**). **d.** Cell-type proportions. **e.** Dot plot of canonical markers and LAE- or SAE-enriched genes derived from bulk RNA-seq (see Fig. 1f). **f.** Violin plots of LAE and SAE markers. **g.** UMAP of re-clustered secretory cells. **h.** Proportions of LAE- or SAE-derived cells across secretory subsets. **i.** Marker expression across secretory subsets. **j.** Heatmap of top 10 markers. **k.** Co-expression of *SCGB1A1*, *SFTPB*, and *MUC5B*. **l.** RNA *in situ* hybridization for secretory markers in SAE. Scale bars, 20 μm.

To further resolve secretory cell heterogeneity, we reclustered secretory and hybrid secretory-ciliated cell populations from both LAE and SAE, identifying five distinct subsets (**Fig. 2g**). While *SCGB1A1*, a shared secretory marker, was broadly expressed across subsets, DASC markers (*SFTPB*, *SCGB3A2*, *RNASE1*) were most highly expressed in a single subset composed of 97% SAE-derived cells after normalization to equal total LAE and SAE cell numbers (**Fig. 2h-j**). We, therefore, designated this subset as the DASC population. The DASC population also expressed *MUC5B* (**Fig. 2i**), a secretory mucin required for maintaining airway innate mucosal defense. scRNA-seq also revealed the heterogeneity of co-expression of *SCGB1A1*, *SFTPB*, and *MUC5B* in SAE secretory cells, whereas LAE secretory cells predominantly expressed *SCGB1A1* and/or *MUC5B* and lacked *SFTPB* (**Fig. 2k**). This heterogeneous expression pattern in SAE secretory cell populations was confirmed by RNA *in situ* hybridization (RNA-ISH) (**Fig. 2l**). Collectively, these data identified DASCs as a distinct, SAE-enriched secretory cell population defining distal airway identity.

### Region- and cell-type specific chromatin regulatory landscapes in human airway epithelia

To define chromatin regulatory elements underlying region- and cell-type-specific transcriptional programs in airway epithelia, we performed single-nucleus (sn) RNA/ATAC-seq on LAE and SAE cultures from three matched donors (**Fig. 3a, Extended Data Fig. 4a**). UMAP analysis of integrated gene expression and chromatin accessibility data identified clusters corresponding to major airway epithelial cell types (**Fig. 3b, Extended Data Fig. 4b-d**), closely mirroring those observed in scRNA-seq data (**Fig. 2**). Notably, DASCs formed a distinct cluster composed predominantly of SAE-derived cells (**Fig. 3c**), characterized by high expression of canonical marker genes (*SFTPB*, *SCGB3A2*, *RNASE1*) (**Fig. 3d, Extended Data Fig. 4e**) and selective chromatin accessibility at DASC marker loci, including *SFTPB* and *RNASE1* (**Fig. 3e, Extended Data Fig. 4f**).

**Fig. 3.**
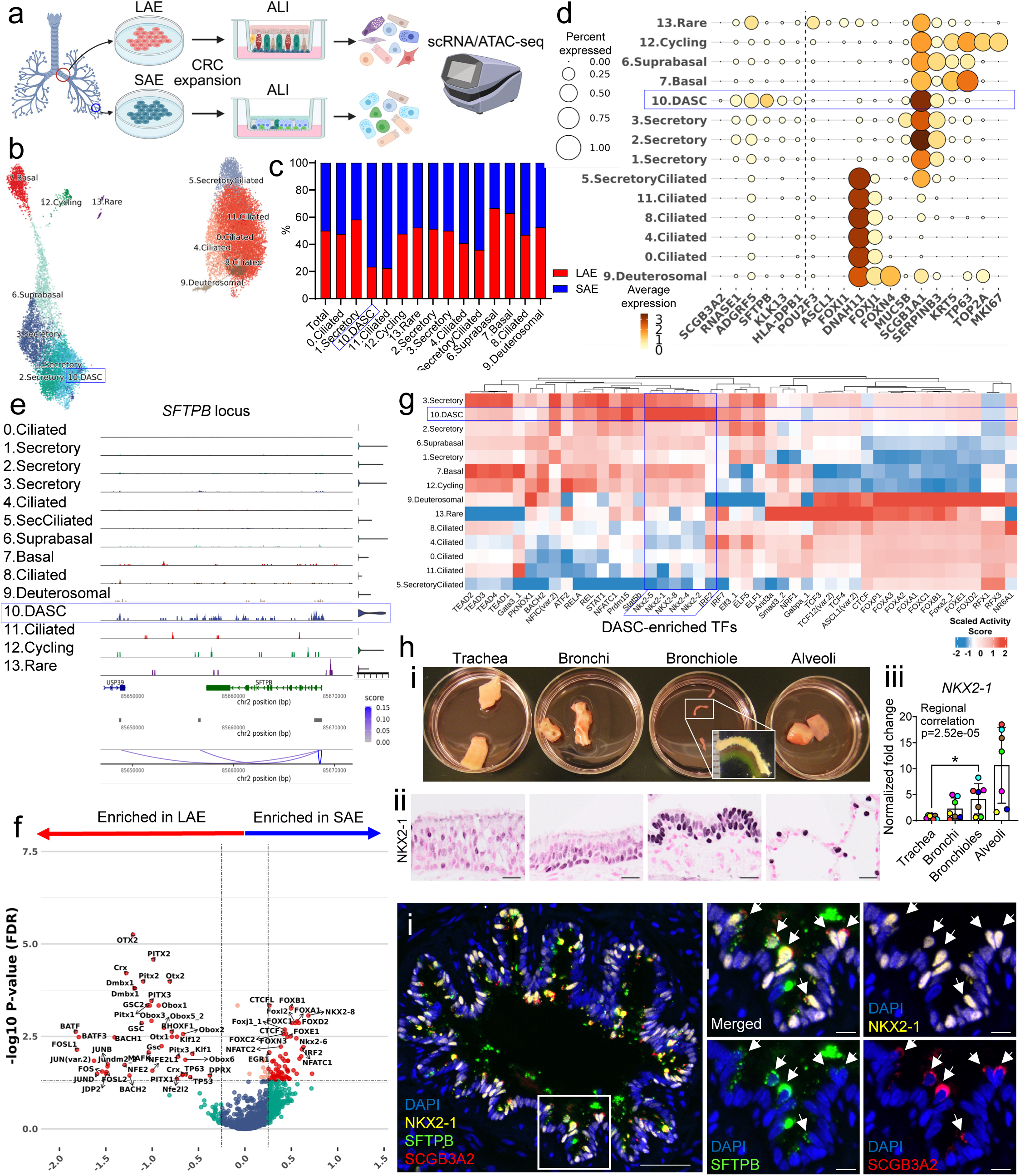
snRNA/ATAC-seq defines region- and cell-type-specific chromatin accessibility in LAE and SAE. **a.** Experimental design. **b.** UMAP of epithelial cells from LAE and SAE (21,005 cells; n = 3). **c.** Cell-type proportions. **d.** Dot plot of epithelial markers. **e.** ATAC-seq peaks at the *SFTPB* locus. **f.** Differentially enriched transcription factor motifs (SAE versus LAE). **g.** Top 10 enriched motifs across cell types. **h.** Proximal-to-distal NKX2-1 gradient in human airways: (**i**) microdissected airway regions; (**ii**) immunohistochemistry (scale bar, 20 μm); and (**iii**) regional transcript expression analyzed by a linear mixed-effects model. *p < 0.05. **i.** Immunofluorescence of NKX2-1 in SFTPB^+^SCGB3A2^+^ DASCs (arrows). Scale bars, 50 μm (10 μm insets).

snRNA/ATAC-seq analysis further revealed region- and cell-type specific open chromatin regions and associated transcription factor (TF) motifs. SAE-enriched TF motifs included members of the *NKX* and *FOX* families (**Fig. 3f**), which are known regulators of distal airway and alveolar specification during lung development^28, 29^. In contrast, LAE-enriched TF motifs included *KLF* and AP-1 family members, including *JUN* and *FOS*, which are associated with epithelial proliferation and barrier remodeling^30, 31^. *TEAD* motifs were preferentially enriched in basal and a subset of secretory cell populations (**Fig. 3g**), consistent with their progenitor properties associated with YAP/TAZ signaling^32^. In contrast, *RFX* family TFs (*RFX1*, *RFX3*), key regulators of motile ciliogenesis^33^, exhibited higher activities in ciliated and deuterosomal cells compared to other cell types. Additionally, *ASCL1*, a canonical regulator of neuroendocrine cell differentiation^34, 35^, was significantly enriched in rare cell populations. DASC-enriched TF motifs included members of *NKX* family (*NKX2-1*, *NKX2-2*, *NKX2-5*) as well as the interferon regulatory factor *IRF2*, compared to other secretory cell subsets, suggesting a specific role for these TFs in maintaining DASC identity and function. Among these DASC-enriched TFs, NKX2-1 exhibited a proximal-to-distal gradient of increasing expression in adult non-diseased human airways, as confirmed by immunohistochemistry and qRT-PCR analysis of microdissected lung tissues from distinct airway regions (**Fig. 3h, Supplementary Table 3**). Furthermore, immunofluorescence (IF) studies identified robust NKX2-1 expression in SFTPB^+^SCGB3A2^+^ DASCs within non-diseased human bronchiolar tissues (**Fig. 3i**), supporting a link between the NKX2-1-mediated transcriptional programs and DASC identity.

### NKX2-1 is required for DASC differentiation in SAE

snATAC/RNA-seq analyses of LAE and SAE cultures identified NKX2-1 as a strong candidate TF to regulate distal airway regional identity and epithelial homeostasis, in part mediated by DASCs. To explore this observation, we employed CRISPR/Cas9 gene editing to inactivate NKX2-1 in primary human LAE and SAE cultures (**Fig. 4a**). Inference of CRISPR Edits (ICE) analysis^36^ demonstrated highly efficient NKX2-1 knockout (KO) in undifferentiated LAE and SAE cells, which was maintained throughout ALI differentiation (**Extended Data Fig. 5a**) and further validated at the protein level by Western blotting (**Fig. 4b, Extended Data Fig. 5b**). Morphologically, SAE cultures exhibited a robust induction of mucin-granule-containing goblet cells following NKX2-1 deletion (**Fig. 4c**). Whole-mount IF studies revealed that NKX2-1 KO SAE exhibited a significant reduction in SFTPB^+^ DASCs accompanied by a proportional increase in MUC5AC^+^ goblet cells, a phenotype largely absent in NKX2-1 KO LAE (**Fig. 4d-e, Extended Data Fig. 5c**). The fractional area of acetylated α-tubulin^+^ signals, reflecting ciliated cell abundance, remained unchanged in both LAE and SAE following NKX2-1 KO (**Fig. 4e**). Importantly, increased MUC5AC expression and reduced DASC marker expression in NKX2-1 KO SAE were evident from the onset of ALI culture and persisted through full differentiation at day 28 (**Extended Data Fig. 5d**), indicating that NKX2-1 loss in SAE progenitors impairs physiological differentiation, most notably DASC identity.

**Fig. 4.**
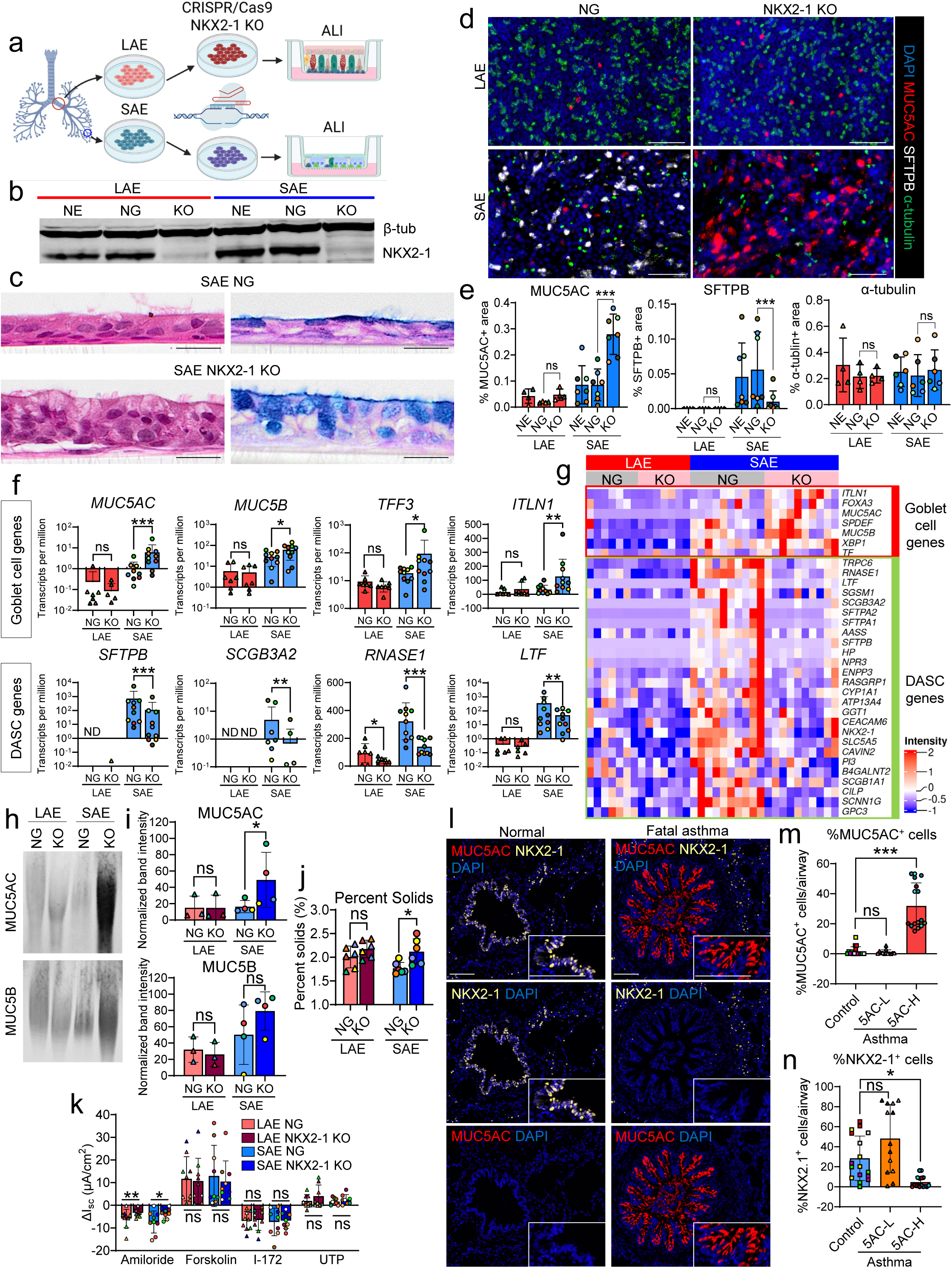
NKX2-1 is required for DASC identity in SAE. **a.** Experimental design. **b.** Western blot of NKX2-1. NE, non-electroporation control; NG, negative gRNA control. **c.** H&E (left) and Alcian blue-periodic acid-Schiff (AB-PAS; right) staining. Scale bar, 20 μm. **d, e.** Whole-mount immunofluorescence of NG and NKX2-1 KO LAE and SAE (**d**) with quantification (LAE, n = 4; SAE, n=7) (**e**). Scale bar, 100 μm. **f.** Bulk RNA-seq of goblet and DASC markers (LAE, n = 7; SAE, n = 10); unadjusted p values are shown for predefined transcripts. ND, not detected. **g.** Heatmap of goblet cell or DASC marker expression. **h, i.** Western blot of MUC5AC and MUC5B in apical secretions (**h**) with quantification (**i**). **j.** Mucus concentration (%solids) (n = 6). **k.** Electrophysiological measurements using Ussing chambers (n = 10). **l-n.** Immunofluorescence of MUC5AC and NKX2-1 in control (n = 5) and fatal asthma bronchioles (**l**) (n = 7), with quantification for frequency of MUC5AC^+^ cells (**m**) and NKX2-1^+^ cells (**n**). Scale bar, 100 μm. ns, not significant; *p < 0.05; **p < 0.01; ***p < 0.001 (linear mixed-effects model).

To further investigate region-specific transcriptional changes induced by NKX2-1 KO, we conducted bulk RNA-seq on NKX2-1 KO or negative gRNA control (NG) LAE and SAE cultures. In NKX2-1 KO SAE, a marked upregulation of goblet cell markers (*MUC5AC*, *MUC5B*, *TFF3*, *ITLN1*) was observed, juxtaposed to downregulation of DASC markers *(SFTPB*, *SCGB3A2*, *RNASE1, LTF*) (**Fig. 4f-g**), consistent with whole-mount IF data (**Fig. 4d-e, Extended Data Fig. 5c**). These changes in transcriptional profiles were relatively unique to SAE NKX2-1 KO and minimally observed in NKX2-1 KO LAE (**Fig. 4f-g, Extended Data Fig. 5e-f**). This pattern was supported by enrichment of pathways associated with goblet cell metaplasia in SAE NKX2-1 KO (**Extended Data Fig. 5g**). Importantly, markers of other airway epithelial cell types, including deuterosomal/ciliated (*FOXN4*, *FOXJ1*), basal (*KRT5*, *TP63*), neuroendocrine (*ASCL1*), tuft (*POU2F3*), and ionocytes (*ASCL3*, *FOXI1*), were not significantly altered by NKX2-1 KO in either LAE or SAE (**Extended Data Fig. 5h**). These data suggest that NKX2-1 is specifically required for DASC differentiation in SAE and that disruption of NKX2-1-dependent transcriptional programs drives a secretory cell fate switch from physiological DASCs toward pathologic, MUC5AC-dominant goblet cells.

To assess the impact of the loss of NKX2-1 on small airway mucus concentration-dependent biophysical properties, we analyzed major secreted mucin production, mucus concentration, and ion transport-mediated airway surface hydration in NKX2-1 KO vs NG LAE and SAE. MUC5AC secretion onto the apical surfaces after 7 days of accumulation was significantly increased in NKX2-1 KO SAE compared with NG SAE, whereas no increase was observed in NKX2-1 KO LAE (**Fig. 4h-i**). MUC5B secretion exhibited a similar upward trend (**Fig. 4h-i**), consistent with a significant increase in *MUC5B* at the transcript level (**Fig. 4f**). Consistent with these findings, NKX2-1 KO SAE exhibited significantly elevated mucus concentration, as indexed by percent solids (**Fig. 4j**). Ussing chamber measurements revealed a significant reduction in amiloride-sensitive short-circuit current, an index of epithelial sodium channel (ENaC)-mediated transepithelial Na^+^ and fluid absorption, in both LAE and SAE following NKX2-1 KO, consistent with downregulation of *SCNN1B* and *SCNN1G* transcripts (**Extended Data Fig. 5h**). Similar reductions in ENaC-mediated fluid absorption have been reported in LAE cultures following IL-13-induced goblet cell metaplasia^37, 38^, suggesting that decreased ENaC activity is associated with transcriptional reprogramming toward a goblet cell differentiation. In contrast, no significant differences in CFTR-mediated Cl^-^ and fluid secretion were detected between NKX2-1 KO and NG LAE or SAE, as assessed by forskolin stimulation and CFTR inhibitor I-172, (**Fig. 4k**), despite a modest reduction in *CFTR* transcript levels in NKX2-1 KO SAE (**Extended Data Fig. 5h**). Similarly, TMEM16A-mediated UTP-stimulated Cl^-^/fluid secretion was unaffected by NKX2-1 deletion. Together, these findings indicate that excessive mucin secretion in NKX2-1 KO SAE exceeded the epithelial capacity to maintain an appropriate airway surface liquid volume, resulting in hyperconcentrated mucus. Consistent with this notion, hypoxia-associated genes (*EGLN3*, *SPAG4*, *P4HA1*)^39^ were significantly upregulated in NKX2-1 KO SAE under conditions in which mucus was not removed for up to three days (**Extended Data Fig. 5h**), indicative of apical epithelial hypoxia associated with a hyperconcentrated thickened mucus layer.

To extend these in vitro findings to human disease, we examined distal airways (bronchioles) from individuals with fatal asthma, characterized by MUC5AC^+^ goblet cell metaplasia^21^, and from previously healthy control subjects (**Supplementary Table 3**). IF analysis of bronchiolar tissues from individuals with fatal asthma revealed reduced NKX2-1 expression accompanied by increased MUC5AC^+^ goblet cells (**Fig. 4l**). Given the substantial inter-airway heterogeneity in goblet cell abundance across asthma bronchioles^21^, bronchioles were stratified into MUC5AC-high and MUC5AC-low groups for morphometric analysis. NKX2-1 expression was significantly reduced in MUC5AC-high bronchioles (**Fig. 4m-n**), revealing an inverse relationship between NKX2-1 and MUC5AC in distal airway epithelia and linking our in vitro findings to a clinically relevant disease context.

Collectively, these studies identify a region-specific role for NKX2-1 in regulating DASC differentiation and mucus homeostasis in the human distal airway epithelium.

### NKX2-1-high basal cells with secretory features define a unique DASC progenitor population

Within human distal airway epithelia, both basal cells and DASCs are reported to serve as progenitors for local epithelial cell lineages^17, 19, 40, 41^. However, the cellular origins of the DASC lineage have not been defined. Given the unique presence of DASCs in SAE in vivo and in vitro, coupled to region-specific regulation of transcriptional programs and chromatin/TF regulatory elements between SAE and LAE basal cells (**Fig. 2, 3, Extended Data Fig. 3h**), we hypothesized that a subset of SAE basal cells are intrinsically programmed for DASC differentiation in vivo and that this program is maintained in vitro.

First, to test whether SAE progenitor cells could function autonomously to generate DASCs, undifferentiated SAE and LAE progenitor cells were mixed in defined ratios (100:0, 75:25, 50:50, 25:75, and 0:100), and SFTPB^+^ areas were quantified post-differentiation as an index of DASC numbers. With proliferation capacities comparable across LAE and SAE progenitors (**Extended Data Fig. 6a-b**), SFTPB^+^ areas positively correlated with the proportion of SAE progenitors (**Extended Data Fig. 6c-e**), indicating that regional lineage specificities were maintained independently of inter-population interactions. qRT-PCR confirmed a statistically significant positive correlation between SAE marker expression (*SFTPB*, *SCGB3A2*) and the proportion of SAE progenitors throughout differentiation (**Extended Data Fig. 6f**), supporting autonomous lineage identity differences between SAE and LAE progenitors.

Next, we used 3D organoid cultures to define differentiation trajectories from purified progenitor populations, including LAE and SAE basal cells and DASCs, isolated from fully differentiated planar LAE and SAE ALI cultures by fluorescence-activated cell sorting (FACS) (**Fig. 5a**). DASCs were labeled using a lentiviral GFP reporter driven by the human *SCGB3A2* promoter (h*SCGB3A2*-GFP)^42, 43^. Promoter specificity to SCGB3A2^+^ DASCs was validated by whole-mount IF (**Fig. 5b, Extended Data Fig. 6g**), FACS (**Fig. 5c**), and qRT-PCR/bulk RNA-seq data (**Extended Data Fig. 7a-c**). Basal cells and DASCs were separately isolated using the basal cell surface marker NGFR and h*SCGB3A2*-driven GFP expression (**Fig. 5c**). FACS resolved two GFP^-^populations from LAE: 1) NGFR^+^ basal cells; and 2) NGFR^-^ non-basal cells. The LAE NGFR^-^GFP^-^population expressed higher levels of *SCGB1A1*, *MUC5B*, and *FOXJ1*, with minimal expression of DASC markers (*SFTPB*, *SCGB3A2*), consistent with non-DASC secretory and ciliated cells (**Extended Data Fig. 7a-b**). In contrast, SAE segregated into four distinct populations based on NGFR and GFP expression (**Fig. 5c**). The SAE NGFR^-^GFP^+^ population was enriched for DASC markers (*SCGB3A2*, *SCGB1A1*, *SFTPB*) and *EGFP*, confirming successful DASC isolation, whereas the SAE NGFR^-^GFP^-^ population expressed *MUC5B* and *FOXJ1* with lower levels of DASC markers and *EGFP*, consistent with non-DASC secretory and ciliated cells (**Extended Data Fig. 7a-b**). NGFR^+^ SAE basal cells also segregated into two subsets, including: 1) NGFR^+^GFP^-^cells expressing high levels of basal cell markers (*KRT5*, *TP63*); and 2) NGFR^+^GFP^+^ cells expressing intermediate levels of DASC markers and *EGFP*, lower than those in NGFR^-^GFP^+^ DASCs. Bulk RNA-seq confirmed these transcriptional patterns, demonstrating that while canonical basal cell markers (*KRT5*, *TP63*) were expressed across all NGFR^+^ basal cell populations but not in DASCs, SAE NGFR^+^GFP^+^ basal cells uniquely co-expressed basal and DASC/secretory transcriptional signatures (**Fig. 5d, Extended Data Fig. 7b**). Both SAE NGFR^+^ basal cell subsets expressed distal airway basal cell markers identified in vivo (*GPC3*, *MFSD2A*, *APOD*)^19, 44^, whereas these transcripts were only weakly expressed in LAE NGFR^+^ basal cells (**Fig. 5d**), indicating preserved distinct region-specific transcriptional identities within the basal cell populations.

**Fig. 5.**
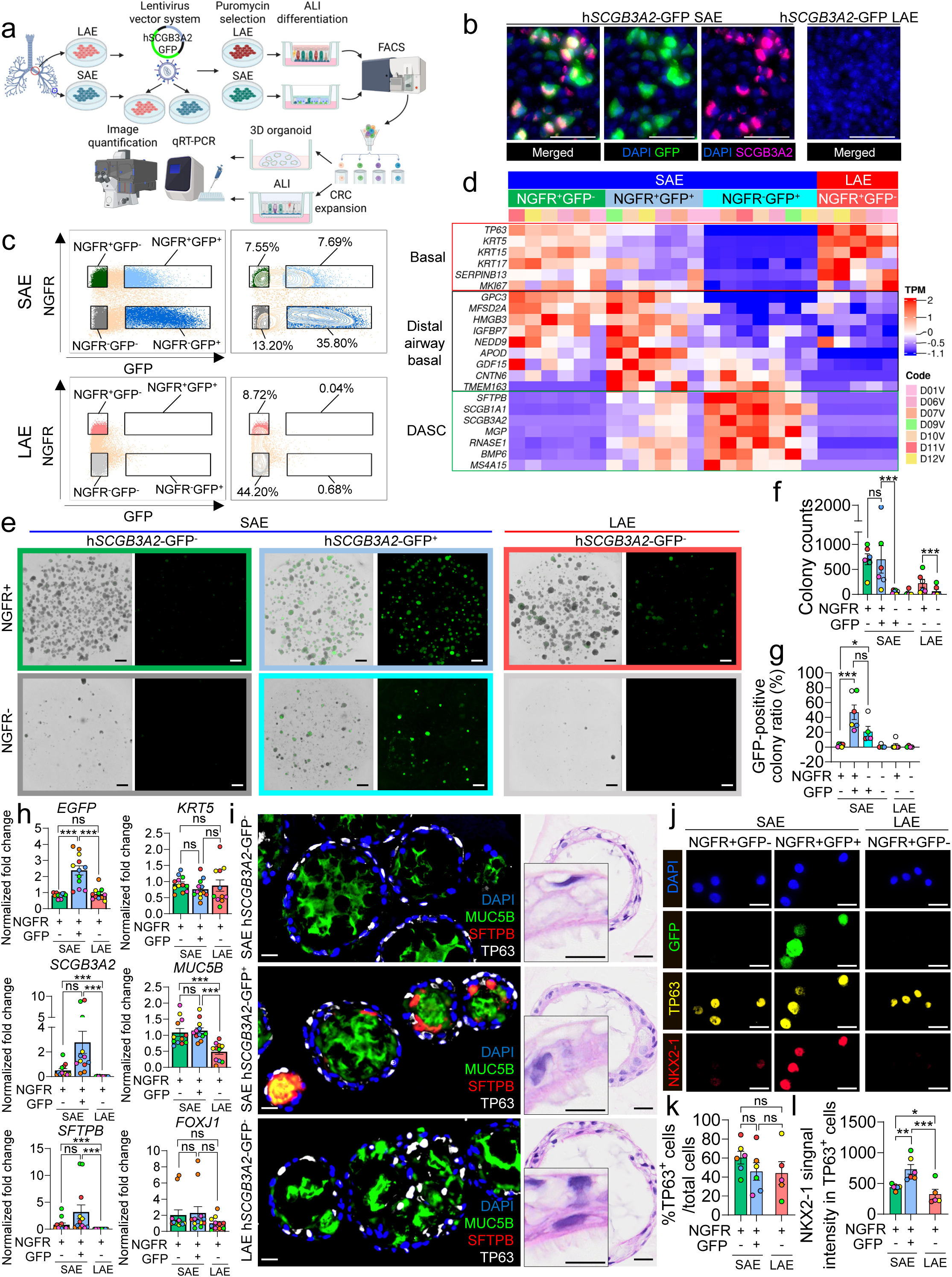
Identification of a distal airway-specific NKX2-1 high DASC progenitor basal cell population. **a.** Experimental design. **b.** Immunofluorescence of h*SCGB3A2*-GFP-transduced LAE and SAE. Scale bar, 100 μm. **c.** FACS isolation using NGFR and h*SCGB3A2*-GFP. **d.** Heatmap of basal, distal basal, and DASC markers in sorted populations. **e-g.** Organoid GFP imaging (**e**), colony quantification (**f**), and GFP^+^ organoid frequency (**g**). Scale bar, 500 μm. **h.** qRT-PCR of organoids derived from sorted basal populations. **i.** Immunofluorescence and H&E staining of organoids from sorted basal cell populations. Scale bars, 20 μm (10 μm insets). **j-l.** Immunofluorescence of sorted NGFR^+^ basal subsets (**j**), TP63^+^ quantification (**k**), and NKX2-1 intensity (**l**). Scale bar, 20 μm. ns, not significant; *p < 0.05; **p < 0.01; ***p < 0.001 (linear mixed-effects model).

In organoid assays, both SAE NGFR^+^ basal cell subsets generated the highest numbers of colonies under PC-S medium conditions, followed by LAE NGFR^+^GFP^-^ basal cells (**Fig. 5e-f**). SAE NGFR^-^GFP^+^ DASCs formed fewer colonies, suggesting a lower proliferative capacity compared to NGFR^+^ basal cell populations, whereas LAE or SAE NGFR^-^GFP^-^ populations generated few colonies, reflecting the absence of progenitor populations in these subsets. Notably, GFP^+^ DASCs were most frequently detected in colonies derived from SAE NGFR⁺GFP⁺ basal cells (∼50%), followed by colonies derived from pre-existing SAE NGFR⁻GFP⁺ DASCs (∼20%), but were rare or absent in colonies derived from LAE or SAE NGFR^+^GFP^-^ basal cells (**Fig. 5e, g**). qRT-PCR analysis of organoids demonstrated expression of *EGFP* and DASC markers (*SFTPB*, *SCGB3A2*) enriched in cultures derived from SAE NGFR^+^GFP^+^ basal cells compared with those derived from LAE or SAE NGFR^+^GFP^-^ basal cells (**Fig. 5h**). Consistently, SFTPB protein was detected almost exclusively in organoids derived from SAE NGFR^+^GFP^+^ basal cells (**Fig. 5i**). In contrast, all NGFR^+^ basal cell populations retained the capacity to generate MUC5B^+^ secretory, ciliated, and TP63^+^ basal cells (**Fig. 5h-i**), indicating shared multipotency for major airway epithelial lineages. Together, these findings indicate that DASCs arise either from pre-existing DASCs or from a specific subset of SAE NGFR⁺ basal cells defined by a hybrid basal-DASC transcriptional profile.

To determine whether the SAE NGFR^+^GFP^+^ basal cell population represents a transient basal-to-DASC intermediate or a stable progenitor state, FACS-isolated NGFR^+^ basal cell subsets were expanded to confluence under CRC conditions, followed by differentiation under ALI conditions. After 28 days, cultures derived from SAE NGFR^+^GFP^+^ basal cells retained higher expression of *GFP* and DASC markers (*SFTPB*, *SCGB3A2*) compared with cultures derived from other basal cell subsets (**Extended Data Fig. 7c**). Consistently, qRT-PCR analysis of CRC-expanded undifferentiated LAE and SAE progenitors isolated directly from fresh human lung tissues revealed higher baseline expression of DASC markers (*SFTPB*, *SCGB3A2*) and DASC-associated TFs (*NKX2-1*, *FOXA2*) in SAE progenitors compared with LAE progenitors, despite comparable expression of basal markers (*KRT5*, *TP63*) (**Extended Data Fig. 7d**). Together, these results support the conclusion that SAE NGFR^+^GFP^+^ basal cells with hybrid basal-DASC features represent a stable, DASC-directed progenitor population rather than a transient intermediate state. Finally, because NKX2-1 is required for DASC differentiation in SAE (**Fig. 4**), we examined whether NKX2-1 is highly expressed in SAE NGFR^+^GFP^+^ basal cell populations. Although the proportion of TP63^+^ basal cells was comparable across FACS-isolated NGFR^+^ basal cell subsets, IF analysis revealed significantly higher NKX2-1 signal intensity per TP63^+^ basal cell (**Fig. 5j-l**), coupled to a trend toward a greater fraction of NKX2-1^+^ cells (**Extended Data Fig. 7e**), in the SAE NGFR^+^GFP^+^ basal population compared to LAE and SAE NGFR^+^GFP^-^ basal populations, consistent with the increased NKX2-1 activity predicted by regulatory network analysis of bulk RNA-seq data (**Extended Data Fig. 7f**). Importantly, the proportion of NKX2-1^+^ basal cells was positively correlated with the frequency of GFP^+^ organoid colonies (**Extended Data Fig. 7g**), consistent with an essential role for NKX2-1-dependent transcriptional programs in DASC differentiation (**Fig. 4**). Together, these data identify an SAE-specific NKX2-1-high basal cell population with hybrid canonical basal-DASC signatures as a dedicated progenitor for DASCs.

### Spatial correlation between NKX2-1-high distal airway basal cells and DASCs in human distal airways in vivo

To link our in vitro findings to human in vivo distal airway epithelial biology, we examined whether the region-specific NKX2-1-high DASC progenitor basal cell subset identified in vitro was present in human distal airway tissues (**Supplementary Table 3**) and whether its regional distribution correlated with that of DASCs. KRT5^+^ basal cells were distributed along the airway axis with a progressive reduction in overall abundance toward distal airways (**Fig. 6a-c**). In contrast, NKX2-1^+^KRT5^+^ basal cells were readily detected in distal bronchiolar epithelia where SFTPB^+^ DASCs were present but were rare in proximal bronchiolar regions lacking DASCs in control human lungs (**Fig. 6a-b**). Quantitatively, the NKX2-1^+^KRT5^+^ basal cell subset comprised approximately 54% and 83% of all KRT5^+^ basal cells in distal bronchiolar (< 1 mm) and terminal bronchiolar (TB) regions, respectively (**Fig. 6d**), indicating that elevated NKX2-1 expression is a defining feature of distal airway basal cells in vivo. Consistent with this distribution, a subset of KRT5^+^ basal cells co-expressing SFTPB, indicative of a basal-DASC hybrid state, was detected in intact bronchioles (1∼2 mm diameter) with increasing frequency toward more distal regions, accounting for ∼18% and ∼36% of KRT5^+^ basal cells in distal bronchiolar (< 1 mm) and TB regions, respectively (**Fig. 6e-f**). The majority of SFTPB^+^KRT5^+^ basal cells also expressed NKX2-1 (**Fig. 6c, Extended Data Fig. 8a**). Similarly, KRT5^+^ basal cells co-expressing SCGB3A2 were also observed in DASC-enriched distal airways (**Extended Data Fig. 8b**). Importantly, the spatial distributions of both NKX2-1^+^KRT5^+^ and SFTPB^+^KRT5^+^ basal cell subsets positively correlated with the abundance of SFTPB^+^KRT5^-^ DASCs along the distal airway axis (**Fig. 6g-i**), supporting a lineage relationship between NKX2-1-high distal airway basal cells, including those exhibiting hybrid basal-DASC features, and DASCs, consistent with our in vitro findings (**Fig. 5, Extended Data Fig. 7**).

**Fig. 6.**
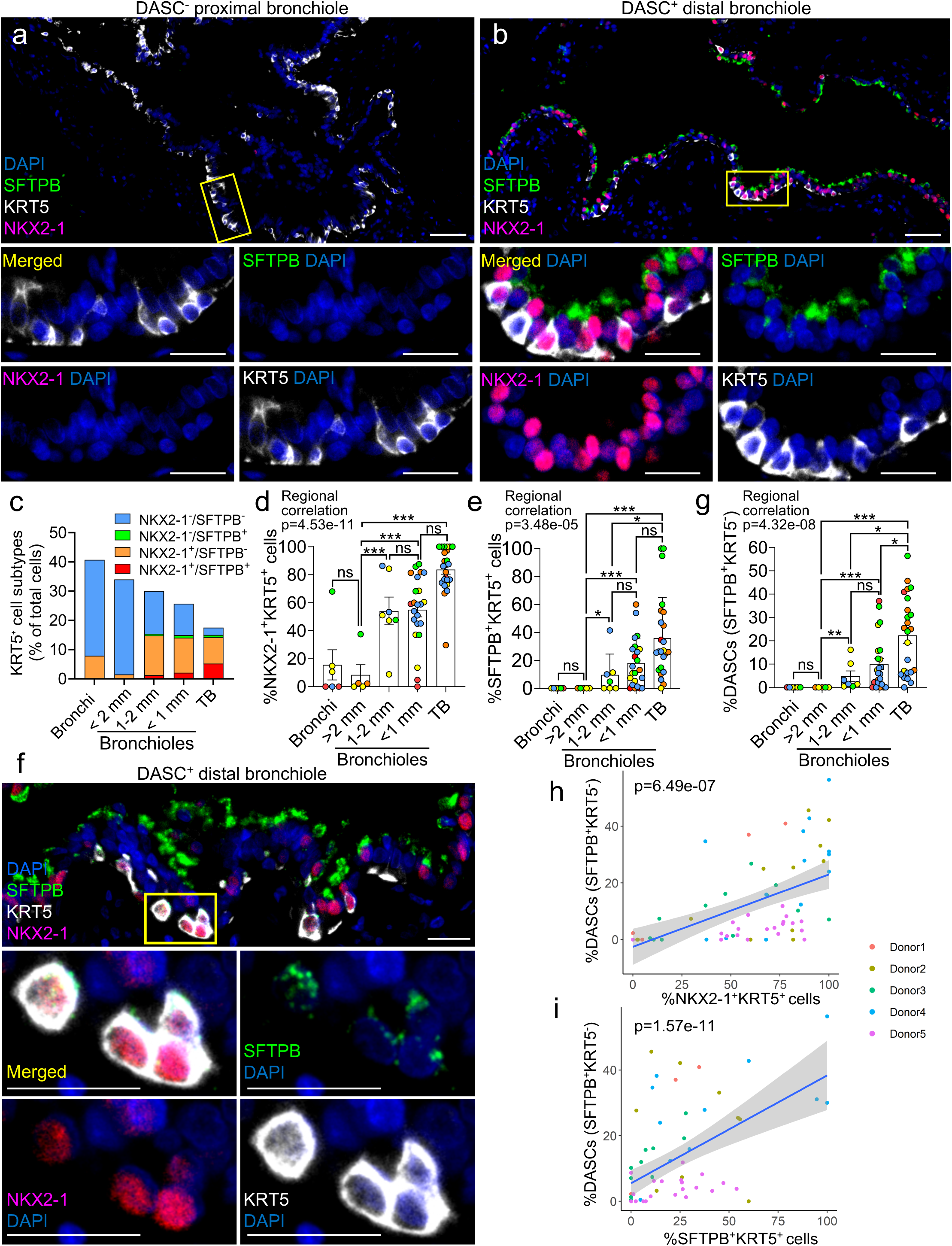
Spatial correlation between NKX2-1 high basal cells and DASCs. **a, b.** Immunofluorescence of selected markers in representative DASC^+^ distal (**a**) and DASC^-^ proximal (**b**) bronchioles. Scale bar, 50 μm (20 μm insets). **c.** Quantification of NKX2-1^+^ and/or SFTPB^+^ cells among KRT5^+^ basal cells across regions. TB, terminal bronchioles. **d, e.** Proportions of NKX2-1^+^KRT5^+^ (**d**) and SFTPB^+^KRT5^+^ (**e**) basal cells among epithelial cells along the proximal-distal axis. **f.** Immunofluorescence of NKX2-1^+^KRT5^+^SFTPB^+^ basal cells in a distal bronchiole. Scale bar, 20 μm. **g.** Frequency of DASCs (SFTPB^+^KRT5^-^) across airway regions. **h, i.** Correlation between DASC abundance and NKX2-1^+^KRT5^+^ (**h**) or SFTPB^+^KRT5^+^ (**i**) basal cells. Each color represents airways from a single donor. n = 5. Regional differences and correlations were analyzed using a linear mixed-effects model. ns, not significant; *p < 0.05; **p < 0.01; ***p < 0.001.

## Discussion

Airways undergo approximately 23 generations of branching until alveolar surfaces are reached, an architecture that efficiently delivers air to the large alveolar surface area within the limited thoracic space. This branching pattern results in an exponential increase in epithelial surface area as airways extend distally, rendering the small/distal airways the dominant airway compartment and a critical site for lung host innate defense. Mucus is a key component of MCC, and small airways must tightly regulate both local and intraregional mucus properties for mucosal defense. Dysregulation of mucus regulatory processes impairs MCC and promotes mucus accumulation in small airway/bronchiolar regions, as observed in MOLDs^3–6, 21, 45^. The absence of backup host defense mechanisms, including submucosal gland secretions^46^ and cough clearance^13^, further contributes to the vulnerability of small/distal airways to muco-obstructive diseases. Reflecting these region-specific host defense requirements, the distal airway epithelium is comprised of specialized cell populations distinct from large airways, including a recently characterized distal airway-specific secretory cell population, i.e., DASCs^17–19^. These cells likely regulate mucin and antimicrobial peptide secretion, cytokine and chemokine production, and transepithelial ion transport, likely dominating small airway host defense and MCC properties. Loss of this cell population and its protective host defense functions are linked to MOLD pathogenesis^18, 21, 22^. Hence, elucidating distal airway/DASC biology is essential to define the mechanisms of distal airway vulnerability and to establish a mechanistic basis for therapies aimed at improving outcomes in MOLDs.

By integrating bulk and single-cell transcriptional profiling of LAE and SAE cultures, we defined region- and cell-type-specific transcriptional programs across the two airway regions. Notably, SAE-enriched genes (*SFTPB*, *SCGB3A2*, *RNASE1*) were predominantly expressed in secretory cell populations, including DASCs, whereas LAE-enriched genes (*KRT13*, *KRT14*) were expressed in basal cell populations (**Fig. 2e-f**), identifying these lineages as principal drivers of regional transcriptional divergence. Consistent with these findings, proteomic analysis of apical secretions demonstrated a more diverse secretome in SAE compared to LAE (**Fig. 1h, Extended Data Fig. 2**). The greater spectrum of host defense proteins secreted by SAE suggests that distal airway epithelia compensate for the absence of submucosal glands through enhanced epithelial secretory function.

During embryonic lung development, lung cell specification and morphogenesis are precisely regulated by integrated transcriptional signaling between mesenchymal and epithelial cells^28, 34, 47^. Among key pulmonary TFs, NKX2-1 is one of the earliest markers of lung epithelial progenitors, distinguishing respiratory from gut lineages within the foregut endoderm, and NKX2-1 is required for subsequent distal lung epithelial specification^28, 29^. Due to its predominantly lung-restricted expression (with additional expression in the thyroid and brain)^48^, NKX2-1 has been widely used to isolate lung progenitor populations derived from human and mouse pluripotent stem cells^49, 50^. Despite extensive evidence supporting its developmental activity, NKX2-1 function in adult human lung epithelium, particularly within conducting airway epithelia, remains less well-defined. Our chromatin accessibility analysis of primary LAE and SAE cultures revealed regionally distinct epithelial chromatin landscapes (**Fig. 3**), consistent with prior studies implicating TFs associated with proximal-distal patterning in adult mouse and human airways, including NKX2-1^51^, FOXA2^52^, SOX9^53^, and GATA6^54^. Amongst these TFs, we identified NKX2-1 as one of the most highly active TFs in DASCs.

Mechanistically, CRISPR/Cas9-mediated NKX2-1 depletion in SAE progenitors resulted in a near-complete loss of DASC phenotypes, whereas LAE remained largely unaffected by this maneuver, providing direct evidence for a region-specific regulatory role of NKX2-1 in human conducting airway epithelia (**Fig. 4**). In parallel, NKX2-1 loss induced pronounced MUC5AC-dominant goblet cell metaplasia in SAE cultures, phenocopying a hallmark pathological feature of small airways commonly observed in MOLDs^18, 21, 22, 55, 56^ (**Fig. 4l-n**). This apparent “anti-mucous” function of NKX2-1 is consistent with prior reports demonstrating its role in suppressing mucous cell metaplasia and Th2-driven inflammation in allergen-challenged mouse models^51, 57^. Notably, the NKX2-1 KO phenotype in SAE contrasts with that in alveolar epithelium, where NKX2-1 loss drives transdifferentiation of AT2 cells into KRT8^+^ stressed transitional cells^58^, rather than goblet cells. Importantly, replacement of DASCs by goblet cells in NKX2-1 KO SAE was accompanied by increased mucus concentration that can markedly impair MCC^59, 60^. Overall, these data suggest a highly integrated, region-specific dual role for NKX2-1 in preserving DASC identity while suppressing goblet cell differentiation, both of which are critical for maintaining effective small airway host defense (**Extended Data Fig. 8c**).

Because in vitro SAE cultures recapitulate key in vivo SAE molecular features in the absence of mesenchymal or immune interactions, these findings suggest that airway epithelial progenitors retain intrinsic, region-specific epigenetic programs. Although DASCs themselves exhibit progenitor capacity to generate multiple airway epithelial lineages^17, 19^, the identity of the upstream progenitor population responsible for DASC specification has not been defined. Using a h*SCGB3A2* promoter-driven GFP reporter, we segregated NGFR⁺ SAE basal cells into two distinct subsets and found that h*SCGB3A2*-GFP⁺ SAE basal cells preferentially generated colonies containing DASCs (**Fig. 5**). Notably, NKX2-1 was highly enriched in this DASC progenitor basal population, consistent with its functional requirement for DASC differentiation and supporting a key regulatory role for NKX2-1 in the DASC lineage. Interestingly, h*SCGB3A2*-GFP⁻ SAE basal cells retained broad progenitor capacity for multiple airway epithelial lineages but failed to generate DASCs, despite expressing elevated levels of distal airway basal cell markers as compared to h*SCGB3A2*-GFP⁻ LAE basal cells (**Fig. 5d-i, Extended Data Fig. 7a-c**). This observation suggests that these h*SCGB3A2*-GFP^-^ SAE basal cells may represent basal populations from proximal bronchiolar regions where DASCs are emerging but not yet predominant, exhibiting intermediate features between large airway and distal airway DASC progenitor basal cells.

Notably, the h*SCGB3A2*-GFP⁺ DASC progenitor basal cell subset was characterized by a hybrid transcriptional profile incorporating both basal and secretory/DASC features (**Fig. 5d**, **Extended Data Fig. 7a-b**). This observation raised the question of whether these cells represent a transient intermediate state en route to DASC differentiation or a phenotypically stable distal airway-specific progenitor population. Our organoid-based comparisons of DASC differentiation capacity across distinct basal cell subsets supported the latter, consistent with prior studies. First, scRNA-seq analyses of freshly excised normal human lung tissue demonstrated that distal airway basal cells exhibited lower expression of canonical basal markers (*TP63*, *KRT5*) and higher expression of secretory/DASC-associated genes (*SFTPB*, *NKX2-1*, *RNASE1*) compared with large airway basal cells^19^, indicating that a basal-secretory hybrid transcriptional state is a stable feature of distal airway basal cells in vivo. Second, secretory-primed basal cell populations have been identified in normal human proximal airway tissues and shown to possess clonogenic and multilineage differentiation capacity following isolation^61^, consistent with the clonal progenitor features observed in our CRC-expanded h*SCGB3A2*-GFP⁺ SAE basal cells (**Extended Data Fig. 7c**). Importantly, expansion of secretory-primed basal cells with aberrant basaloid features has been reported in distal lung tissues from patients with IPF^61^, juxtaposed to evidence of physiological DASC loss in IPF lungs^22^, suggesting that dysregulated differentiation within this progenitor population contributes to pathological epithelial remodeling. Third, SCGB3A2⁺ airway progenitor cells have been identified in early fetal human lungs (8-11 weeks post-conception) and shown to differentiate into multiple epithelial lineages^34^, supporting the existence of a developmentally conserved progenitor state with secretory features. Finally, our in vivo IF analysis confirmed the presence of an NKX2-1-high basal cell population with DASC features in normal human distal airways and demonstrated its spatial correlation with DASCs (**Fig. 6**). Collectively, these findings suggest a distal airway-specific basal cell population with stable hybrid basal-DASC features as a progenitor for DASC differentiation.

In conclusion, we establish a spatially resolved human in vitro molecular atlas of airway epithelia that defines regulatory programs governing region-specific epithelial identity and host defense. This framework delineates transcriptional programs required to maintain physiological distal airway epithelial cell composition and function, exemplified by NKX2-1. We further identify a mechanistic link between disrupted NKX2-1-dependent differentiation programs and impaired distal airway mucus properties, highly relevant to the pathogenesis of MOLDs. In addition, we define a specific lineage relationship between an NKX2-1-high distal airway basal cell population and DASCs. Collectively, these findings establish a foundation for interrogating region-specific airway epithelial biology and physiology in the human distal airways, a compartment that is difficult to access in vivo.

## Methods

### Human Subjects

De-identified excess surgical pathology tissues were obtained from the University of North Carolina (UNC) Tissue Procurement and Cell Culture Core under protocol #03-1396 approved by the UNC Biomedical Institutional Review Board (IRB). Written informed consent was obtained from tissue donors or their legally authorized representatives in accordance with institutional and federal guidelines. Cystic fibrosis (CF) lung tissues were procured from donors undergoing lung transplantation. Control lungs from individuals without prior chronic lung disease, deemed unsuitable for transplantation, as well as lungs from subjects with fatal asthma, were obtained through Carolina Donor Services (Durham, NC), the National Disease Research Interchange (Philadelphia, PA), or the International Institute for Advancement of Medicine (Edison, NJ). Excised specimens were processed immediately for primary epithelial cell isolation or RNA extraction, or fixed in 10% neutral buffered formalin (Fisher Chemical, SF100-4) for 72 h prior to paraffin embedding to generate formalin-fixed paraffin-embedded (FFPE) tissue sections^62, 63^. In total, 67 donor lungs were included in this study: previously healthy individuals (n = 60) and subjects with fatal asthma (n = 7). Among these, 57 lungs from previously healthy donors were used to establish matched large airway epithelial (LAE) and small airway epithelial (SAE) cultures. Donor demographic information is provided in **Supplementary Tables 1** and **3**. LAE and SAE cells were expanded and differentiated under identical culture conditions for all comparative analyses.

### LAE and SAE cell isolation, expansion, and differentiation

Freshly excised human lung tissues were processed under sterile conditions in a laminar flow biosafety cabinet. LAE cells were isolated from cartilaginous airways (trachea and bronchi > 2 mm in diameter) by enzymatic digestion with freshly prepared pronase (10 mg/mL, Roche, 10165921001) and Accutase (5 mL, MilliporeSigma, A6964) for 2 h at room temperature (RT) on an orbital shaker, as previously described^15, 64^.

SAE cells were obtained using two complementary approaches: microdissection or bulk enzymatic digestion. For microdissection, bronchioles were identified within distal lung tissue by the absence of cartilage and an outer diameter < 2 mm. The absence of cartilage and submucosal glands was confirmed by histological assessment. Bronchioles were dissected free from surrounding parenchyma under light microscopy, immediately placed in F12 medium on ice, longitudinally opened, and subjected to enzymatic digestion with pronase (10 mg/mL) and Accutase (5 mL) for 2 h at RT^15^. Enzymatic activity was neutralized with 10% fetal bovine serum (FBS) (GeminiBio, S12450), and cells were collected by centrifugation (600 × g, 2 min, 4°C). For bulk digestion, distal lung tissue (∼3 × 3 cm) was excised from peripheral regions beyond bronchi > 2 mm in diameter, and visceral pleura was removed. Tissue was minced and digested with pronase (10 mg/mL) and Accutase (5 mL) for 2 h at RT with agitation. Following addition of 10% FBS, the suspension was filtered through a 40-µm cell strainer (pluriSelect, 43-50040-01) and centrifuged (600 × g, 2 min, 4°C). If erythrocyte contamination was present, red blood cell lysis buffer (Invitrogen, 00-4333-57) was applied for 2 min, followed by neutralization with DMEM (Gibco, 11995065) containing 10% FBS. Cells were centrifuged and resuspended for culture.

Isolated LAE and SAE cells were expanded using a conditional reprogramming culture (CRC) technique^15, 25^. Cells were co-cultured with mitomycin C-treated 3T3-Swiss albino (ATCC, CCL-92) feeder cells on type1 collagen (Advanced BioMatrix, 5005)-coated tissue culture dishes (Corning, 430167) in DMEM supplemented with 10 µM Y-27632 (Enzo Life Sciences, ALX-270-333-M025). For selected samples used in mass spectrometry-based secretome analysis (see **Supplementary Table 2**), cells were expanded using Bronchial Epithelial Cell Growth Medium (BEGM) without Y-27632 or feeder cells. At 70-90% confluence, cells were either passaged or cryopreserved in the cryopreservation media (NIPPON Genetics, CS-02-002/BB05) for subsequent use. Passage 2 (P2) LAE and SAE cells were seeded onto human placental type IV collagen (MilliporeSigma, C5533)-coated 0.4-µm pore size Millicell inserts (Sigma-Aldrich, PIHP01250) or equivalent Transwell inserts (Corning, 3460) at ∼5.0 × 10^5 cells/cm². Cells were maintained in CRC medium for 24 h, then transitioned to differentiation media (UNC air-liquid interface [ALI], PneumaCult-ALI (STEMCELL Technologies, 05001), or PneumaCult-ALI-S (STEMCELL Technologies, 05050) (see **Supplementary Table 2**). Upon reaching confluence, apical medium was removed to establish an air-liquid interface (ALI), and differentiation medium was supplied to the basal compartment only. Basal medium was replaced twice weekly, and apical surfaces were washed weekly with PBS (Gibco, 14190144). After 28 days of differentiation at ALI, LAE and SAE cultures were used for downstream experimental analyses. For experiments using cryopreserved cells, vials were rapidly thawed in a 37°C water bath, washed with PBS, and expanded on collagen-coated tissue culture dishes in CRC medium prior to differentiation.

### Mixed LAE and SAE cell cultures

LAE and SAE cells isolated from matched human large and small airway tissues were expanded using the CRC method as described above. Upon confluence, expanded LAE and SAE cells were combined at defined ratios (100:0, 75:25, 50:50, 25:75, and 0:100), seeded onto Transwell inserts, and differentiated under ALI conditions for 28 days. Following full differentiation, cells were harvested for total RNA extraction and qRT-PCR analysis or fixed in 4% paraformaldehyde for whole mount immunofluorescence.

### H&E and AB-PAS histology analysis of LAE and SAE cultures

LAE and SAE culture inserts were fixed in 4% paraformaldehyde (PFA) (Electron Microscopy Science, 15710), paraffin-embedded, and sectioned at 5 µm. Sections were stained with hematoxylin and eosin (H&E) or Alcian blue-periodic acid-Schiff (AB-PAS) and imaged using a VS200 slide scanner [Olympus (Evident Scientific)].

### Alveolar type 2 cell isolation, expansion, and differentiation

Alveolar epithelial type 2 (AT2) cells were isolated from the peripheral regions of human lung tissue, followed by purification of HT2-280 (Terrace Biotech, TB-27AHT2-280) positive AT2 cells using magnetic-activated cell sorting (MACS) (Miltenyl Biotech, 130-091-051), as previously described^26^. Human AT2 cells were initially cultured in serum-free, feeder-free (SFFF) medium, and alveolosphere cultures were passaged after 10-14 days. Passage 2 or 3 AT2 cells were then seeded onto Transwell inserts coated with Collagen III (Advanced BioMatrix, 5021) and Biolaminin 332 (BioLamina, LN332-0502), following the Stemcell Technologies protocol (https://www.stemcell.com/technical-resources/how-to-generate-alveolar-monolayers-from-atii-organoids.html#more) and cultured in SFFF medium supplemented with Y-27632 and maintained under submerged conditions for 2 days. Once confluence was achieved, cultures were air-lifted, the medium was changed to SFFF, and cells were maintained under ALI conditions for an additional 5 days.

### Lentiviral production of human SCGB3A2-reporter vector

A 600-bp fragment corresponding to the human *SCGB3A2* promoter region (chr5: 147878111-147878710)^34, 43^ was synthesized (Azenta) and cloned into the lentiviral backbone p-Lenti CMV GFP Puro (Addgene plasmid #658-5) by replacing the CMV promoter using ClaI (New England Biolabs, R0197S) and XbaI (Thermo Scientific, FD0864) restriction sites. All constructs were verified by Sanger sequence prior to viral production. Lentivirus was produced in HEK293T cells using a third-generation packaging system. HEK293T cells (Takara Bio, 632180) were expanded in 150-mm tissue culture dishes (Corning, 430599) in DMEM supplemented with 10% FBS and 1% penicillin-streptomycin until ∼80% confluence. For transfection, 10 µg of transfer plasmid encoding the human SCGB3A2 promoter, 5 µg of pCMV-VSV-G, and 8 µg of psPAX2 were diluted in 1 mL Opti-MEM (Gibco, 31985070). Separately, 35 µL of transfection reagent (Sigma-Aldrich, 6366546001) was diluted in 1 mL Opti-MEM. The DNA and transfection reagent solutions were combined, gently mixed, and incubated at RT for 20-30 min to allow complex formation. The mixture was then added dropwise to HEK293T cells. After 6 h, transfection medium was replaced with fresh medium. Viral supernatant was collected ∼60 h post-transfection, clarified by centrifugation at 4°C (4,000 rpm, 10 min), and filtered through a 0.45-µm low protein-binding membrane filter. Viral titer (Takara Bio, 631281) was assessed, and viral supernatants were used immediately or stored at −80°C until use.

#### Lentivirus vector-mediated gene transfer to LAE and SAE cultures

Primary LAE and SAE cells were plated on collagen-coated tissue culture plastic dishes in CRC medium and maintained at 37°C in a humidified incubator with 5% CO_2_. At ∼50% confluence, culture medium was removed and replaced with 2.5 ml lentiviral supernatant supplemented with 2 µg/ml polybrene (Millipore Sigma, TR-1003-G) and diluted 1:1 with CRC medium. Cells were incubated for 6 h at 37°C. Following transduction, cultures were washed twice with PBS and replenished with fresh CRC medium. Upon reaching confluence, cells were dissociated using Accutase and seeded onto collagen-coated tissue culture plastic dishes with feeder cells under CRC medium supplemented with puromycin (2 μg/mL, Gibco, A1113802). Cells were passaged to Transwell or Millicell inserts for differentiation under ALI conditions as described above. After differentiation, cultures were harvested for flow cytometric analysis and fluorescence-activated cell sorting (FACS).

### Organoid culture of sorted LAE and SAE cells

Primary LAE and SAE cells were transduced with the h*SCGB3A2*-EGFP plasmid and cultured under ALI conditions using PneumaCult ALI-S medium supplemented with puromycin for 28 days to full differentiation. On day 28, dissociated cells were stained with LIVE/DEAD Fixable Near-IR (Invitrogen, L34975), followed by Fc receptor blocking using Human TruStain FcX (BioLegend, 422302). Cells were labeled with BV421-conjugated anti-human CD271 (NGFR) antibody (BD, 562562) and sorted using a FACSAria2 cell sorter (BD). Sorted cells were resuspended in DMEM supplemented with 10% FBS and mixed 1:1 with Growth Factor Reduced Matrigel (Corning, 354230). A total of 5 × 10³ live cells per Matrigel dome were plated into 24-well plates (VWR, 10062-896). Cells were maintained in expansion medium^65^ for the first 3 days at 37°C in a humidified incubator with 5% CO_2_, followed by differentiation using PneumaCult ALI-S medium. The medium was replaced every 3 days. On day 14 of differentiation, images were obtained for counting colony numbers using a BZ-series all-in-one microscope (KEYENCE). Colony numbers were manually counted from bright field or fluorescence images acquired under identical conditions across different groups. The percentage of EGFP-positive colonies was determined based on GFP signal intensity adjusted for appropriate threshold identical across different groups. Cultures were also harvested for the following downstream analyses. For histological analysis, Matrigels were embedded in 1% low-melting-point agarose (RPI, A20070), fixed in 4% PFA for 30 min at RT, and washed with PBS. For RNA extraction, Matrigels were digested with TrypLE Express (Gibco, 12563029) for 5 min at 37 °C, followed by neutralization with DMEM with 10% FBS.

### Colony-forming assay of LAE and SAE progenitor cells

For colony-forming assays (**Extended Data Fig. 6a-b**), 1 × 10³ CRC-expanded LAE or SAE cells were resuspended in 50 µL UNC ALI medium and mixed 1:1 with growth factor-reduced Matrigel (Corning). The cell-Matrigel mixture (100 µL total) was seeded onto 6.5-mm Transwell inserts (Corning, 3422) as previously described^66^. Bronchospheres were cultured for 14-18 days and stained with Calcein AM (1 ng/mL in ALI medium; Invitrogen, C3100MP) prior to imaging. Fluorescent images were acquired using an Olympus IX-81 inverted wide-field microscope [Olympus (Evident Scientific)] with a 2× objective. Colony-forming efficiency (CFE) was calculated as the percentage of spheres formed relative to the number of cells initially seeded.

### Immunofluorescence and immunohistochemistry

Immunostaining was performed on paraffin-embedded human lung tissue sections. Slides were baked at 60°C for 2-4 h, deparaffinized in xylene (2 × 5 min) (epredia, 6601), and rehydrated through graded ethanol (100% 2 × 5 min, 95% 1 × 5 min, 70% 1 × 5 min). Antigen retrieval was performed in 0.1 M sodium citrate buffer (pH 6.0) (MilliporeSigma, C9999) using microwave heating (100% power for 6.5 min, followed by 60% power for 6 min twice) or the steamer (Hamilton Beach, 37530A) for 20 min. Slides were allowed to cool to RT and rinsed in distilled water. For chromogenic immunohistochemistry, endogenous peroxidase activity was quenched with 0.5% hydrogen peroxide (MilliporeSigma, H1009) in methanol (MilliporeSigma, 34860) for 15 min. Slides were washed in PBS and blocked with Blocking One Histo (Nacalai USA, 06349-64) for 1 h at RT. Primary antibodies were diluted in 5% Blocking One in PBS containing 0.1% Triton X-100 (PBS-TX) and incubated overnight at 4°C. Species-specific γ-globulin or serum were used as an isotype control at matched concentrations. After washing in PBS-TX, sections were incubated with species-specific secondary antibodies for 60 min at RT. For fluorescent staining, sections were treated with the Vector® TrueVIEW Autofluorescence Quenching Kit (Vector Laboratories, SP-8400) according to the manufacturer’s instructions and mounted with 4′,6-diamidino-2-phenylindole (DAPI) containing mounting media (Vector Laboratories, H-1800). For 3-3’-Diaminobenzidine (DAB)-based chromogenic detection, slides were incubated with avidin–biotin–peroxidase complex (Vector Laboratories, PK-6100), followed by visualization with DAB (Thermo Scientific Chemicals, H54000.14) and counterstaining with Nuclear Fast Red (MilliporeSigma, N8002) followed by mounting with DPX Mounting Medium (Electron Microscopy Science, 13512). Slides were cover slipped and imaged using a VS200 whole-slide scanner. Antibodies used in this study are listed in **Supplementary Table 4**.

### Whole-mount immunostaining and imaging

Well-differentiated LAE and SAE cultures were washed with PBS and fixed in 4% PFA for an hour at RT. Fixed cells were permeabilized, blocked, and incubated in primary antibodies over night at 4°C as described above^15^. Specific-secondary antibody mixture containing DAPI (Invitrogen, D1306) was used, and images were obtained using a VS200 whole-slide scanner. Quantification of target protein signal was performed using Fiji (**Fig. 4e**) or QuPath (**Extended Data Fig. 6e**). The area positive for signal above a predefined intensity threshold was measured and normalized to total epithelial area.

### RNA in situ hybridization (RNA-ISH)

RNA-ISH was performed on paraffin-embedded 5-µm tissue sections using the RNAscope 2.5 HD Reagent Kit, RNAscope 2.5 HD Duplex Reagent Kit, or RNAscope Multiplex Fluorescent Assay v2 (Advanced Cell Diagnostics, 322350, 322430, or 323100, respectively) according to the manufacturer’s protocols. Slides were deparaffinized in xylene (2 × 5 min) and 100% ethanol (2 × 1 min), followed by hydrogen peroxide treatment for 10 min. Target retrieval was performed in boiling water for 15 min, and sections were incubated with Protease Plus for 15 min at 40°C. Slides were hybridized with custom-designed probes at 40°C for 2 h, and signal amplification and detection were carried out according to the manufacturer’s instructions. Brightfield or fluorescent images were acquired using a VS200 whole-slide scanner.

### RNA isolation and quantitative RT-PCR (TaqMan assays)

For gene expression analysis of freshly dissected lung tissues (**Fig. 3hiii**), surface epithelial cells were isolated from normal trachea and bronchi obtained from previously healthy transplant donors by gentle scraping with a convex scalpel blade into F12 medium, excluding submucosal glands. Cells were pelleted by centrifugation (450 × g, 5 min, 4°C) and resuspended in 1 mL TRI Reagent (MilliporeSigma, T9424). Microdissected small airways and peripheral lung parenchyma were homogenized directly in 1 mL TRI Reagent using a tissue homogenizer (Bertin Technologies). Lysates were clarified by centrifugation, and the supernatant was used for RNA extraction. For cultured cells, undifferentiated CRC-expanded LAE and SAE cells grown on plastic, as well as differentiated LAE and SAE cultures maintained at ALI on Transwell or Millicell membranes, were harvested for RNA isolation using the RNeasy Mini Kit (QIAGEN, 74104) according to the manufacturer’s instructions. For FACS-sorted cells and dissociated cells from Matrigel-organoid cultures, total RNA was extracted using the RNeasy Micro Kit (QIAGEN, 74004). RNA quantity and purity were assessed using a NanoDrop 8000 spectrophotometer (Thermo Fisher Scientific). For cDNA synthesis, 1 µg of total RNA from tissue and ALI cultures and 50 ng total RNA from sorted cells and organoid cultures were reverse transcribed using iScript™ Reverse Transcription Supermix (BIO-RAD, 1708840) at 46°C for 60 min. Quantitative real-time PCR (qRT–PCR) was performed using TaqMan gene expression assays (Applied Biosystems) with SsoAdvanced Universal Probes Supermix (BIO-RAD, 1725280) on a QuantStudio 6 Real-Time PCR System (Applied Biosystems). Gene expression was normalized to housekeeping genes (*GAPDH* or *TBP*) using the ΔΔCt method. Primer and probe information is provided in **Supplementary Table 5**.

### CRISPR/Cas9-mediated gene editing of NKX2-1

*NKX2-1* was disrupted in primary LAE and SAE cells using CRISPR/Cas9 ribonucleoprotein (RNP) electroporation as previously described^67^. Briefly, RNP complexes were assembled by combining synthetic sgRNA (360 pmol; Synthego) targeting the sequence 5′-CGCCGTACCAGGACACCATG-3′ with Cas9-NLS nuclease (80 pmol; Synthego) in 60 µL Buffer R (Invitrogen, MPK10096). Undifferentiated passage 1 or 2 (P1/P2) LAE and SAE cells expanded under CRC conditions were dissociated with Accutase, and 6 × 10⁵ cells were resuspended in 160 µL Buffer R. RNP complexes were mixed with the cell suspension and electroporated using the Neon Transfection System (Invitrogen) at 1600 V for 20 ms (single pulse). Following electroporation, cells were immediately transferred to CRC culture conditions for expansion. Upon reaching confluence, edited cells were dissociated and seeded onto type-IV collagen-coated Millicell or Transwell inserts for differentiation at ALI as described above. After 28 days of ALI differentiation, edited cultures were used for downstream analyses. Editing efficiency was evaluated by isolating genomic DNA (QIAGEN, 69504), PCR amplification of the sgRNA-targeted locus, and Sanger sequencing of amplicons. Indel frequencies were quantified using Synthego Inference of CRISPR Edits (ICE) analysis^36^ (**Extended Data Fig. 5a**). Non-targeting sgRNA sequences 5’-GCACUACCAGAGCUAACUCA-3’ lacking predicted genomic targets were used as negative controls.

### Western blotting assay for MUC5AC and MUC5B

Apical secretions accumulated over 7 days from well-differentiated LAE and SAE cultures were collected by washing the apical surface with 200 µL PBS. Mucin Western blotting assay was performed as previously described^63^. Samples were solubilized in 6 M urea, reduced with 10 mM dithiothreitol (90 min at 37°C), and alkylated with 25 mM iodoacetamide (30 min at RT in the dark). Equal volumes of reduced samples (30 µL) were resolved on 1.2% agarose gels (80 V, 90 min) and transferred to nitrocellulose membranes by vacuum blotting in 4× sodium citrate buffer for 2 h. Membranes were blocked in blocking buffer (LI-COR Biosciences, 927-60001) and incubated with primary antibodies against MUC5B (rabbit polyclonal, H-300, 1:2000, SantaCruz, sc-20119) or MUC5AC (mouse monoclonal, 45M1, 1:2000, Invitrogen, MA5-12178) diluted in blocking buffer supplemented with 0.1% Tween-20. After washing, membranes were incubated with IRDye 680LT donkey anti-rabbit or anti-mouse secondary antibodies (LI-COR, 1:10,000). Signals were detected and quantified using the Odyssey infrared imaging system (LI-COR), with densitometric values normalized to background intensity.

### Western blotting assay

Primary human AT2 cells (positive control) and fully differentiated LAE, and SAE cultures at ALI day 28 were used for protein analysis. Cultures were washed once with 400 μL PBS and incubated for 15 min at 37 °C. Cells were lysed with 100 μL ice-cold RIPA buffer supplemented with 1% SDS (Millipore Sigma, 71736) and 1× Halt protease and phosphatase inhibitor cocktail (Invitrogen, 78440). Lysates were incubated on ice for 30 min with brief vortexing (10 s) every 10 min, followed by centrifugation at 18,000 × g for 15 min at 4 °C. Supernatants were collected and stored at −80 °C until analysis. Protein concentrations were determined using a bicinchoninic acid (BCA) protein assay kit (Thermo Scientific, 23225) according to the manufacturer’s instructions. Equal amounts of protein (40 μg) were mixed with Laemmli sample buffer containing 2-mercaptoethanol (BIO-RAD, 1610710), heated at 60 °C for 30 min, and separated by SDS-PAGE using 4-20% precast TGX gels (BIO-RAD, 4561094). Proteins were transferred onto polyvinylidene difluoride membranes (BIO-RAD) using a wet tank blotting system according to the manufacturer’s protocol (BIO-RAD). Membranes were washed three times with PBST and blocked with blocking buffer for 30 min at RT. Primary antibodies were diluted in fresh blocking buffer and incubated with membranes overnight at 4 °C with gentle agitation. The following day, membranes were washed three times with PBST and incubated with Fluorescence dye-conjugated secondary antibodies (1:10,000) for 60 min at RT. After additional PBST washes, immunoreactive bands were detected using the Odyssey infrared imaging system (LI-COR).

### Bioelectric measurements in Ussing Chamber

Transepithelial ion transport was assessed by short-circuit current (Isc) measurements in differentiated LAE and SAE cultures as previously described^15, 25^. ALI cultures differentiated for 28 days were mounted in Ussing chambers (Physiologic Instruments). Both apical and basolateral compartments (5 mL each) were filled with Krebs-bicarbonate-Ringer solution containing (in mM): 140 Na⁺, 120 Cl⁻, 5.2 K⁺, 1.2 Ca²⁺, 1.2 Mg²⁺, 2.4 HPO₄²⁻, 0.4 H₂PO₄⁻, 25 HCO₃⁻, and 5 glucose. Solutions were maintained at 36 ± 1°C, continuously gassed with 95% O₂/5% CO₂, and buffered to pH 7.4. To measure CFTR-dependent currents, amiloride (MilliporeSigma, A7410) (100 µM, apical) was first applied to inhibit epithelial sodium channels. CFTR was stimulated by bilateral addition of forskolin (MilliporeSigma, F6886) (10 µM), and CFTR-specific current was subsequently inhibited with CFTR inh-172 (MilliporeSigma, C2992) (10 µM, apical). Calcium-activated chloride channel activity was then assessed by apical addition of UTP (Cytiva, 27-2086-01) (100 µM) in the continued presence of amiloride and CFTR inh-172. Changes in short-circuit current (ΔIsc) were calculated as the difference between baseline current immediately prior to agonist or inhibitor addition and the peak response following acute treatment.

### Measurement of mucus concentration

Apical secretions accumulated over 7 days from well-differentiated NKX2-1 knock out (KO) or control LAE and SAE ALI cultures were collected by washing the apical surface with 100 µL PBS. The mucus samples were weighed on a pre-weighed piece of aluminum foil. The samples were heated at 80°C in an oven overnight to allow the liquid content to evaporate completely. The final weight of the dried foil and mucus sample was determined and solid weight % (%solids) was calculated^68, 69^.

### Mass spectrometry analysis of airway epithelial secretome

Apical secretions (100 µL) collected from LAE and SAE cultures were diluted 1:1 with 8 M guanidine hydrochloride (GuHCl). Samples were reduced with 10 mM dithiothreitol (1 h at 65 °C), alkylated with 20 mM iodoacetamide (1 h at RT in the dark), and digested overnight with sequencing-grade trypsin. Following digestion, high-molecular weight glycopeptides were removed by centrifugation through 10-kDa molecular weight cutoff filters (MilliporeSigma, Amicon® Ultra Centrifugal filters, UFC5010). Filtrates were lyophilized and resuspended in 0.1% formic acid prior to analysis. Peptides were analyzed by liquid chromatography-tandem mass spectrometry (LC-MS/MS) using a Q Exactive mass spectrometer (Thermo Fisher Scientific) as previously described^70^. Proteins were identified by database searching against the current human reference proteome and quantified by label-free analysis using Scaffold v4.4.8 (Proteome Software). Quantification was based on normalized total precursor intensity, incorporating unique peptides with ≥95% probability as determined by the Scaffold Local FDR algorithm.

Label-free quantities (LFQ) from mass spectrometry analysis were preprocessed with Bioconductor R package, *MsCoreUtils*. LFQ values were log_2_ transformed, which converted 0s to missing values, and proteins with >50% missing among samples in any treatment groups were excluded from downstream analysis. The log_2_(LFQ) values were normalized by median centering among all samples. Missing values were imputed by quantile regression imputation of left-censored data (QRILC) method, which generated left-censored missing data using random draws from a truncated distribution with parameters estimated using quantile regression. The normalized and imputed log_2_(LFQ) values were used for principal component analyses (PCA), and differential expression analysis using linear mixed-effect models with the dream function from the Bioconductor R package *variancePartition*^71^.

SFTPB LFQ data from four (DD66M, DD70M, DD28N, and DD73N) out of the eighteen donor-derived microdissected SAE cultures and their matched LAE cultures were previously reported^63^ (**Supplementary Table 1**).

### Bulk RNA-sequencing Analysis and Visualization

RNA sequence libraries were prepared from total RNA isolated from well-differentiated LAE and SAE ALI cultures, NKX2-1 KO and control LAE and SAE cultures, and FACS-isolated epithelial cell populations by polyA capture and non-directional library preparation, followed by paired-end 150 bp bulk RNA-sequencing using NovaSeq 6000 platform (Illumina). Raw FASTQ sequence reads were aligned to the human reference genome (GRCh38) utilizing STAR aligner v2.7.10a^72^ with current GENCODE gene annotations^73^. Aligned BAM files were assembled and quantified using StringTie v2^74^, and gene expression values were exported as counts and transcripts per million (TPMs). PCA and hierarchical clustering were performed using TPM values. Counts were normalized using the voom method and differential expression (DE) analysis was performed using the dream function from the Bioconductor R package, *variancePartition*^71^. Genes were included in DE analysis if the total count across all samples exceeded 100 prior to normalization. Heatmaps visualizing differentially expressed genes (DEGs) were generated from TPM values using the Bioconductor R package ComplexHeatmap^75^. Hierarchical clustering was performed using hclust, with distances calculated using the ‘Manhattan’ method implemented in the amap R package^76^. Gene set enrichment analyses (GSEA) was performed using the fgsea^77^ Bioconductor R package with genes pre-ranked by log_2_ fold change). Pathway analyses were conducted using Reactome (v92)^78^ gene sets, along with curated gene sets derived from published studies^15, 39, 79–83^. NKX2-1 transcriptional activity was inferred with decoupleR^84^, using the CollecTRI regulon resource^85^.

### scRNA-seq of LAE and SAE cultures

The apical surface of differentiated LAE and SAE ALI cultures was washed twice with PBS (5 min each, 37 °C) to remove mucus. Cells were dissociated by adding Accutase (500 µL to the apical compartment and 1 mL to the basolateral compartment) and incubated at 37 °C for 10 min until detachment. The apical compartment was gently pipetted to dislodge cells, which were collected and combined with an additional apical wash using Accutase. The cell suspension was neutralized with FBS, centrifuged (300 × g, 2 min, 4 °C), and resuspended in PBS without Ca^2+^ or Mg^2+^ supplemented with 0.01% ultrapure BSA (Invitrogen, AM2616). Cells were filtered through a 40 µm strainer, washed, and resuspended in PBS with 0.01% BSA prior to cell counting and viability assessment. Dissociated cells were processed using the Chromium Controller (10x Genomics) to generate single-cell gel bead-in-emulsions (GEMs), and libraries were prepared using the Chromium Next GEM Single Cell 3′ Kit v3.1 (10x Genomics) according to the manufacturer’s instructions. Following reverse transcription and cDNA amplification, libraries were constructed and purified using SPRIselect reagent (Beckman Coulter, B23317). Sequencing libraries were quantified using TapeStation (Agilent) and sequenced on a NovaSeq 6000 (Illumina) using paired-end reads to achieve a minimum depth of 20,000 read pairs per cell.

Cell Ranger v.6.1.1 (10x Genomics) was used to map raw sequences to the GRCh38 human reference genome, which were annotated based on the GENCODE v37 protein coding gene annotation [gencodegenes.org]. The resulting raw gene and cell count matrices were imported into R v.4.4.1 [r-project.org] using the R package *Seurat* v.5.1.0^1^. *SoupX* and *scDblFinder* were used to remove ambient RNA and doublet cells from the raw count matrices before downstream processing^2,3^. Count matrices for each sample were subsequently filtered to include cells with gene counts between 300 and 7000 that contained fewer than 20% of reads derived from mitochondrial genes. All datasets were integrated using the default canonical correlation analysis (CCA) method implemented by Seurat. All clustering, marker gene identification, and dimensionality reduction were performed using the default parameters in Seurat. Of the 19 identified clusters, four exhibited markedly low numbers of detected genes and total transcript counts and were therefore classified as low-quality clusters and excluded from downstream analyses (5. Low Quality Ciliated, 7. Low Quality Suprabasal, 10. Low Quality Ciliated, and 15. Low Quality Ciliated; **Extended Data Fig. 3**). Remaining clusters were annotated based on the expression of established marker genes corresponding to basal, ciliated, FOXN4^+^ ciliated, cycling, ionocyte/neuroendocrine (NE), secretory, secretory-ciliated, and suprabasal cell populations (**Extended Data Fig. 3a**). Subclusters were subsequently consolidated into major cell types, including basal, cycling, suprabasal, secretory, secretory-ciliated, ciliated, and rare cell populations, for downstream analysis. To compare relative cellular composition between LAE and SAE cultures, cell proportions were normalized to adjusted counts assuming equal total cell numbers from each culture (**Fig. 3h**, **Extended Data Fig. 3d**). Differential gene expression analysis was performed using the hurdle method from *MAST* v.1.30.0, which is a separate R package available from Bioconductor [Bioconductor.org]^4^.

Cell populations annotated as either secretory or secretory-ciliated cells were extracted from the original integrated dataset to identify subtypes of secretory cells. The extracted data were normalized and scaled according to the SCTransform Seurat function before performing dimensionality reduction by PCA in Seurat. Batch effects between samples were corrected by *Harmony* using default parameters with the calculated PCA selected as the input reduction^5^. Uniform manifold approximation and projection (UMAP) analysis based on the corrected PCA was used to display subclusters of secretory cell populations, and DASC cells were annotated by confirming expression of *SFTPB*, *SCGB3A2*, and *RNASE1*. Unless otherwise stated, all Seurat functions were run using the default parameters.

### Combined snRNA-seq and snATAC-seq of LAE and SAE cultures

Following dissociation of differentiated LAE and SAE ALI cultures, nuclei were isolated using the Chromium Nuclei Isolation Kit (10x Genomics) according to the manufacturer’s instructions. Briefly, cells were lysed on ice, and nuclei were purified using a column-based isolation step, followed by debris removal and washing. Purified nuclei were resuspended in nuclei resuspension buffer and counted. Isolated nuclei were loaded onto a Chromium Controller (10x Genomics) to generate single-nucleus GEMs, and libraries were prepared using the Chromium Next GEM Single Cell Multiome ATAC + Gene Expression kit (10x Genomics) according to the manufacturer’s instructions. This workflow enables simultaneous profiling of chromatin accessibility (ATAC) and gene expression from the same nucleus through transposition, barcoding, reverse transcription, and library amplification. Sequencing libraries were quantified using TapeStation (Agilent) and sequenced on a NovaSeq 6000 (Illumina) using paired-end reads to achieve a minimum depth of 25,000 and 20,000 read pairs per nucleus for ATAC and gene expression (RNA) libraries, respectively.

Cell Ranger ARC v.2.0.1 (10x Genomics) was used to process and align raw sequences from each pair of snATAC and snRNA libraries to the GRCh38 human reference genome, which were annotated based on the Gencode v45 protein coding gene annotation [gencodegenes.org]. The resulting raw count matrices for both the snATAC and snRNA libraries were imported into R v.4.4.1 [r-project.org] using either *Signac* or *Seurat*, respectively^1,6^. Processing of snRNA and snATAC count matrices were performed separately prior to integration. *SoupX* and *scDblFinder* were used to conduct ambient RNA and doublet cell removal within snRNA count matrices for each sample, which were also excluded from the final fragment by cell matrices in each snATAC dataset^2,3^. The nucleosome signal and transcription start site (TSS) enrichment scores were calculated by Signac for each snATAC dataset, and cells were included in the final snRNA and snATAC count matrices if the following criteria were met: RNA counts between 1000 and 25000, ATAC counts fewer than 1×10^5^, at least 200 features per cell, at least 5 cells per feature, nucleosome signal no greater than 2.5, TSS enrichment score no less than 2, and no more than 20% of reads from mitochondrial genes. Peak calling was completed using Signac for all cells remaining in the fragment by cell matrix after filtering. The filtered gene expression matrix was normalized and dimensionality reduction calculated by SCTransform and PCA as above. Conversely, the ATAC peak matrix was normalized by the Signac implementation of term frequency inverse document frequency (TFIDF), and dimensionality reduction was calculated by singular value decomposition (SVD) using Signac.

The normalized gene expression and peak matrices for each sample were merged separately before integration and downstream analysis of the two modalities. Additionally, the annotated peaks for each sample were unified using Signac to obtain a harmonized peak list prior to merging each dataset. All data were integrated using Seurat CCA followed by Harmony batch correction^5^. Dimensionality reduction accounting for both the gene expression and peak counts was conducted by weighted nearest neighbors (WNN) analysis and UMAP in *Seurat*^7^. Cell types were annotated based on clustering performed by Seurat and by using established marker genes for specific cell types as above. To compare relative cellular composition between LAE and SAE cultures, cell proportions were normalized to adjusted counts assuming equal total cell numbers from each culture (**Fig. 3c**). Activity scores for transcription factor motifs identified by Signac were calculated using the *chromVAR* package for subsequent differential accessibility analysis^8^. Both differential gene expression analysis and differential accessibility analysis were completed by *MAST* as above, and false discovery rate (FDR) adjusted p-values less than 0.05 were used to determine statistical significance^4^. All source code used for processing and analysis of both the scRNA-seq and snRNA-seq/snATAC-seq data is freely available as a standalone R package on Github [https://github.com/cschasestevens/WASP/].

### Statistical analysis

No statistical methods were used to predetermine sample size. Experiments were not randomized, and investigators were not blinded to group allocation during experiments or outcome assessment, except for immunohistochemistry and immunofluorescence quantification in human lung tissues and FACS-isolated cells as described below, where analyses were performed in a blinded manner. For comparison of gene and protein expression, organoid colony numbers, frequencies of NKX2-1^+^ cells across FACS-isolated basal cell populations, and IHC/IF-positive staining areas, linear mixed-effect models (LMMs) were used with the R package *lme4*^86^, with treatment as a fixed effect and sample donor as a random-effect. Post hoc multiple comparisons were conducted using Tukey-adjusted contrasts implemented in the *multicomp* package^87^. For correlation analyses between gene and protein expression in human lung tissues, across airway regions, or among varying LAE:SAE progenitor ratios, LMMs were fitted using the R lmerTest package^88^ with Satterthwarte’s approximation for effective degrees of freedom. Where appropriate, generalized linear mixed-effects model logistic regression was applied using Poisson distributions for count data. Spearman’s rank correlation coefficients were used for nonparametric correlation analyses. For bulk RNA-seq data, differential gene expression analysis and gene set enrichment analysis were performed using FDR-adjusted p-values to define significance. For targeted comparisons of predefined genes across conditions, unadjusted p values are reported. All experiments were performed with at least three independent biological replicates, with technical replicates (up to quadruplicates). The number of biological replicates (n) for each experiment is specified in the corresponding figure legends. Unless otherwise stated, statistical significance was defined as p < 0.05.

### Quantification of NKX2-1 signal intensity in FACS-isolated cell populations

After FACS, cytospin slides were prepared and fixed with 10% NBF for 30 min. Slides were washed three times with PBS and stored in 100% ethanol at −20 °C until assay. After rehydration, slides were subjected to antigen retrieval by boiling in pH 6.0 buffer for 20 min, followed by blocking with Blocking One Histo for 30 min. Cells were permeabilized by incubation with PBS-TX for 15 min. Primary antibodies were prepared in 1:10 diluted blocking buffer with PBST and applied overnight at 4 °C. On the following day, slides were washed three times with PBS-TX and incubated with secondary antibody mixtures with DAPI for 1 hour at RT. Slides were then washed three times with PBST, incubated with quenching buffer for 2 min, rinsed twice with PBS, and mounted with coverslips. Images were acquired using the VS200 slide scanner under identical settings across all slides.

Scanned images were imported into QuPath (version 0.5.1; Ref https://doi.org/10.1038/s41598-017-17204-5). To increase purity of sorted cells, a fixed nuclear threshold for TP63 recognition was set within each donor, and TP63-positive cells were extracted using the *Positive Cell Detection* function^89^. For each detected TP63-positive cell, the mean

NKX2-1 signal intensity within the nuclear (DAPI-positive) area was quantified. Average intensity in each donor was used for statistical analysis.

### Quantification of KRT5^+^, NKX2-1^+^, and SFTPB^+^ cells in human lung tissue sections

Immunofluorescence images of bronchus and distal lung tissues from five donors were scanned using the VS200 slide scanner. Donor information is listed in **Supplementary Table 3**. Airways were categorized as follows: bronchi with cartilage or submucosal glands (SMG), and airways without cartilage or SMG, which were further classified by basement membrane-based diameter into >2 mm, 1-2 mm, <1 mm, and terminal bronchioles (TB). TBs were defined as airways that opened directly into alveolar ducts or alveoli^63^.

To minimize sampling bias, manual cell counts were performed by investigators blinded to slide annotations, with a maximum of 200 cells per airway. When fewer than 200 cells were present, all cells within the airway were counted. Brightness and contrast thresholds were adjusted to confirm specific staining of NKX2-1 and SFTPB in type II alveolar epithelial cells as a positive control, ensuring removal of nonspecific signals. Cells were then categorized into eight groups based on the positivity or negativity for KRT5, NKX2-1, and SFTPB. The proportion of each category was calculated for all detected airways.

### Quantification of NKX2-1^+^ and MUC5AC^+^ cells in human lung sections obtained from previously healthy individuals and fatal asthma subjects

MUC5CAC^+^ and NKX2-1^+^ cells were quantified in lung sections obtained from previously healthy individuals and subjects with fatal asthma. Donor information is listed in **Supplementary Table 3**. Cell detection was performed based on nuclear DAPI staining. MUC5AC^+^ or NKX2-1^+^ cells were identified using the *Positive Cell Detection* algorithm implemented in QuPath v0.5.1^89^. For each airway, the proportion of MUC5AC^+^ or NKX2-1^+^ cells was calculated relative to the total number of bronchiolar epithelial cells. Asthma airways classified as MUC5AC-high were defined as airways with a proportion of MUC5AC^+^ cells greater than two standard deviations above the mean of the control airways (>7.71% MUC5AC^+^ cells), consistent with prior criteria^21^. For proportional measurements of IF-based morphometric data, logit transformation was performed as p/(1-p) with a small offset of 0.01 or 0.001 to avoid log or division of zeros. Transformed data were analyzed using LMMs implemented in the R packages, lme4 and lmerTest, incorporating appropriate fixed effect variables such as sex and age. To account for repeated measurements within donors, donor identity was included as a random intercept term in LMMs.

## Materials availability

Human SCGB3A2-EGFP plasmid construct will be available upon request.

## Data availability

All read-level bulk, scRNA-seq, and snRNA/ATAC-seq data from this study will be publicly available on the publication date of this manuscript. All other data supporting the findings of this study are available from the corresponding author upon reasonable request.

## Code availability

All codes and scripts used to analyze the scRNA-seq and snRNA/ATAC-seq data included in this manuscript are available from the corresponding author upon reasonable request.

## Acknowledgements

The authors appreciate UNC Marsico Lung Institute Tissue Procurement and Cell Culture Core for providing human lung tissues, and Mariam Lam, Shannon M. Kelly, and Samuel C. Gallant for their contributions to maintaining SAE cultures for protocol optimization. The authors also thank UNC Animal Histopathology Core for histology sectioning services, Janet Dow and UNC Flow Cytometry Core Facility for performing FACS, and High-Throughput Sequencing Facility for performing scRNA-seq and snRNA/ATAC-seq. The FACS experiment was supported in part by P30 CA016086 Cancer Center Core Support Grant to the UNC Lineberger Comprehensive Cancer Center.

This research was supported by grants from National Institutes of Health [NHLBI: R01HL163602 (to K.O.), P01HL164320 (to R.C.B.), R01HL146557 (to P.R.T), R01HL160939 (to P.R.T), R01HL153375 (to P.R.T), U54HL165443 (to K.O., J.S.H, and G.S.P.), T32HL007106 (to S.A.S.), 5T32HL160494-04 (to T.W.); NIDDK: P30DK065988 (to UNC Tissue Core, R.C.B. and M.G.); NIAID T32 AI007062 (to S.A.S.); UM1TR004406 (S.A.S.), K12TR004416 (to S.A.S.)], ALA/AAAAI Allergic Respiratory Disease Award (S.A.S.), Cystic Fibrosis Foundation Research Grant [KATO22G0 (to T.K.), BOUCHE19XX0 (to R.C.B. and K.O.), BOUCHER22G0 (to R.C.B. and K.O.), OKUDA25G0 (to K.O.), MURANO25F0 (to H.Mu.), STEVEN24F0 (to N.C.S.), FURUSH25F0 (to M.F.), CFF RDP ESTHER24R0 (to UNC Tissue Core and M.G.)] Research grant from Cystic Fibrosis Research Institute (to K.O.).

## Author contributions

K.O. conceived and supervised the study, designed and performed experiments, analyzed and interpreted data, coordinated the research team, generated figures, and wrote the manuscript. H.Mu. and N.C.S. performed experiments, analyzed and interpreted data, generated figures, and contributed to manuscript writing. H.Ma., J.S.H., G.S.P., W.K.O., S.H.R., P.R.T., and R.C.B. provided conceptual input, technical expertise, and critical revisions of the manuscript. A.B.W. and L.F. optimized protocols for SAE cell isolation and culture. S.N. and A.M.C. performed bulk RNA-seq, scRNA-seq, and snRNA-/ATAC-seq experiments. N.C.S., N.M., and H.D. analyzed sequencing data and generated associated figures. G.C., Z.Y.W., R.E.L., L.S., A.S., and S.D. performed in vitro LAE and SAE cultures, qRT-PCR, Western blotting, and lentiviral production. R.C.G. performed RNA *in situ* hybridization. M.C. provided technical guidance in microscopy. L.C.M. contributed to plasmid cloning. E.M. and M.F. performed immunohistochemistry on human lung tissues and quantified distal airway basal cell populations. R.M.I. and K.H. provided technical assistance with FACS experiments and data analysis. B.R. and M.K. performed mass spectrometry of LAE and SAE apical secretions and analyzed the data. N.L.Q., D.M.C., S.E.B., and M.G. conducted electrophysiological experiments using Ussing chambers and analyzed the data. M.I.G. and B.B. quantified LAE and SAE mucus concentrations. A.L. provided technical support for Western blotting of MUC5AC and MUC5B. S.A.S. performed morphometric analyses of MUC5AC and NKX2-1 expression in control and fatal asthma tissues. T.K., T.A., and Y.M. optimized CRISPR/Cas9-mediated NKX2-1 knockout protocols. T.W. and P.R.T. contributed to interpretation of NKX2-1-mediated chromatin regulation in SAE cultures. G.C. contributed to data interpretation and provided expertise on region-specific transcriptional regulatory mechanisms.

## Extended Data Figure Legends

**Extended Data Fig. 1.**
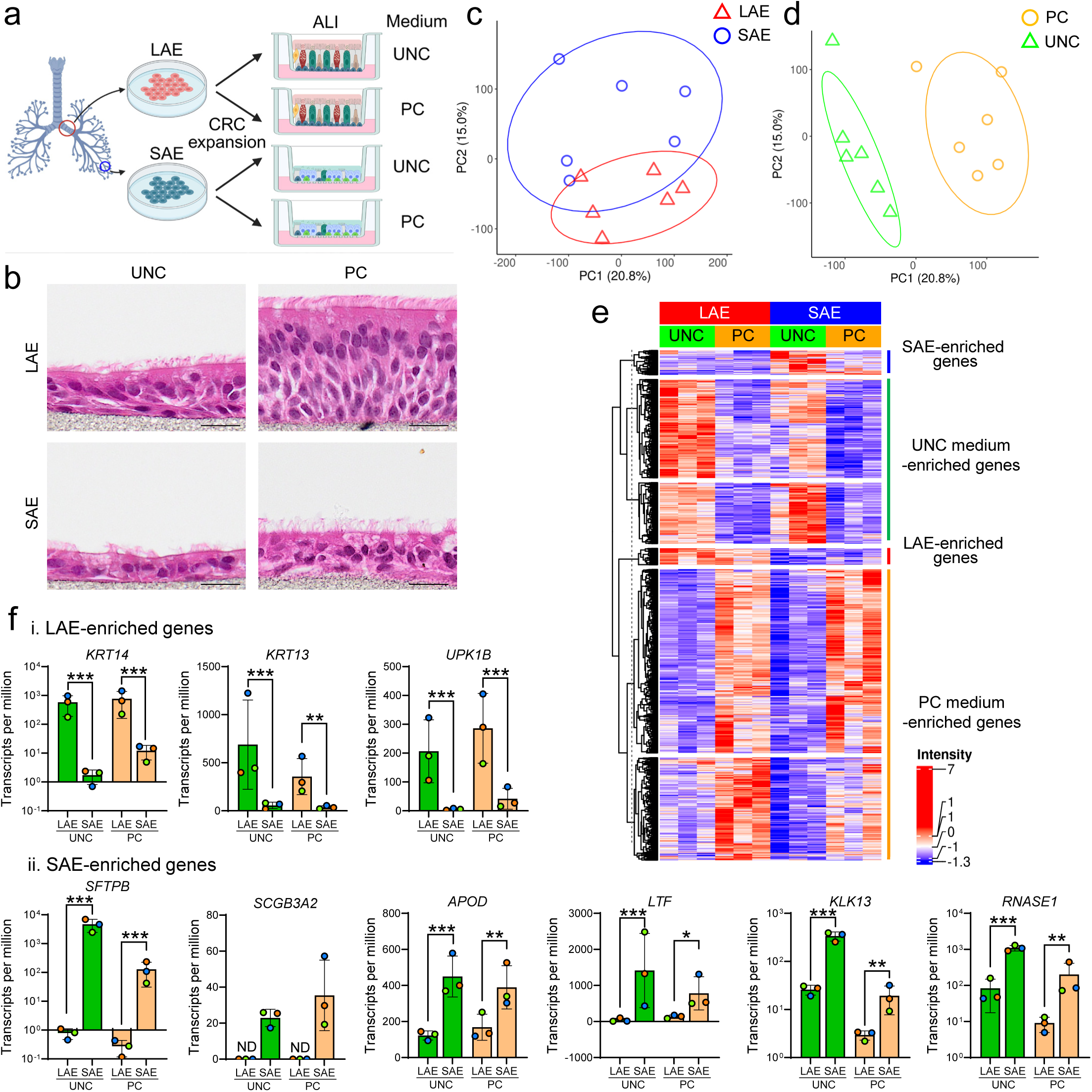
Transcriptional characterization of LAE and SAE cultures under distinct media conditions. **a.** Experimental design (n = 3). UNC, UNC ALI medium; PC, PneumaCult ALI medium. **b.** H&E staining of LAE and SAE ALI cultures. Scale bar, 20 μm. **c, d.** Principal component analysis of LAE and SAE cultures, colored by regional origin (**c**) or culture medium (**d**). **e.** Differentially expressed genes (DEGs) between LAE and SAE and between UNC ALI and PC conditions. **f.** Bulk RNA-seq expression of selected transcripts in LAE and SAE; unadjusted p values are shown for predefined transcripts. ND, not detected. *p < 0.05; **p < 0.01; ***p < 0.001 (linear mixed-effects model).

**Extended Data Fig. 2.**
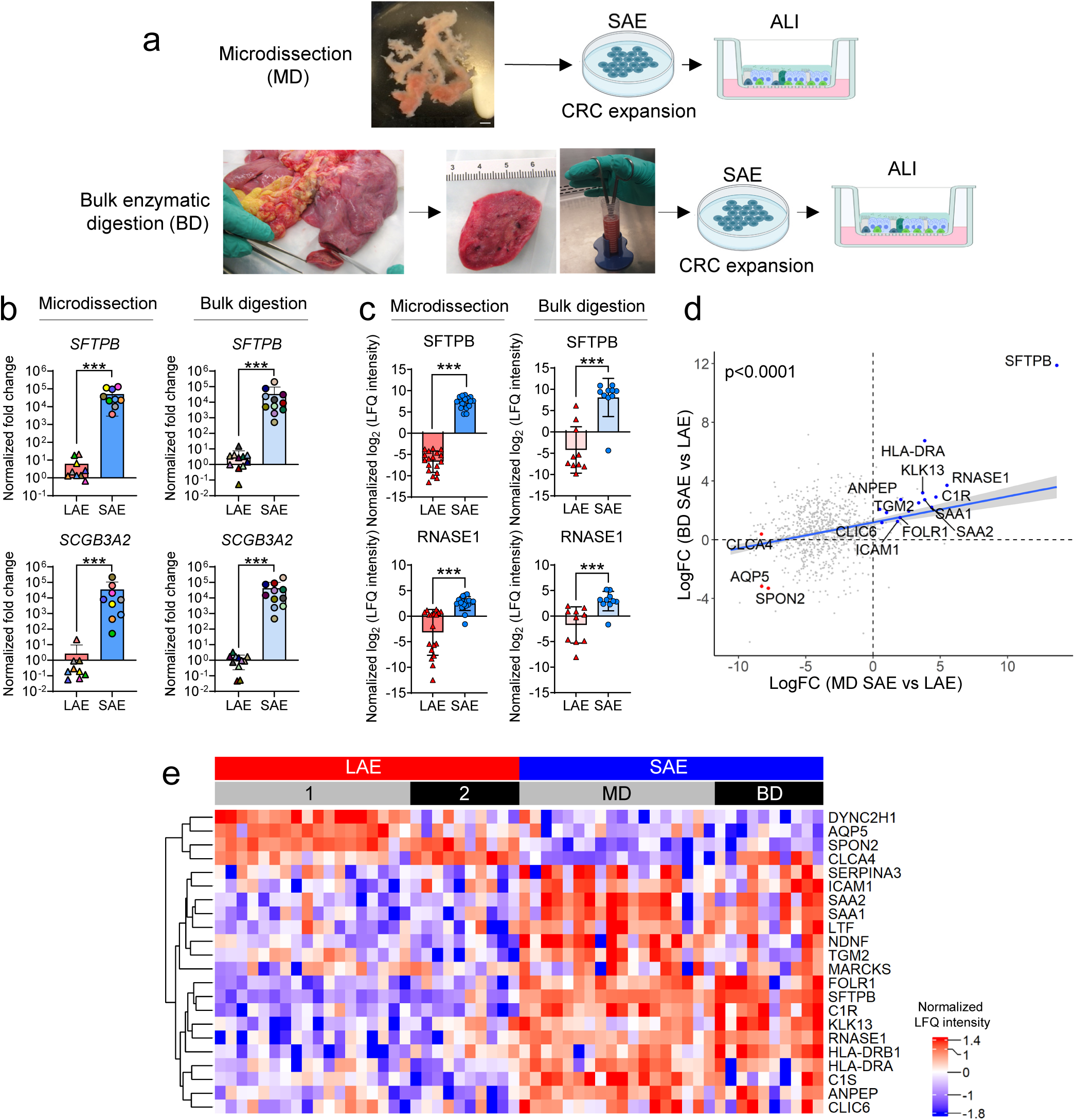
Comparison of SAE isolated by microdissection or bulk digestion. **a.** Schematic of SAE cell isolation by microdissection (MD) or bulk digestion (BD). **b.** Transcript expression in LAE and SAE (LAE vs MD-SAE, n = 9; LAE vs BD-SAE, n = 11). **c.** Mass spectrometry-based label-free quantification (LFQ) of apical secretions from matched LAE and SAE (MD, n = 18; BD, n = 10). **d.** Correlation of fold changes for enriched secretory proteins (SAE vs LAE) between MD- and BD-derived cultures. **e.** Regionally enriched secretory proteins in LAE or SAE secretions. LAE batch 1 and 2 correspond to MD- and BD-derived SAE samples, respectively. ***p < 0.001 (linear mixed-effects model).

**Extended Data Fig. 3.**
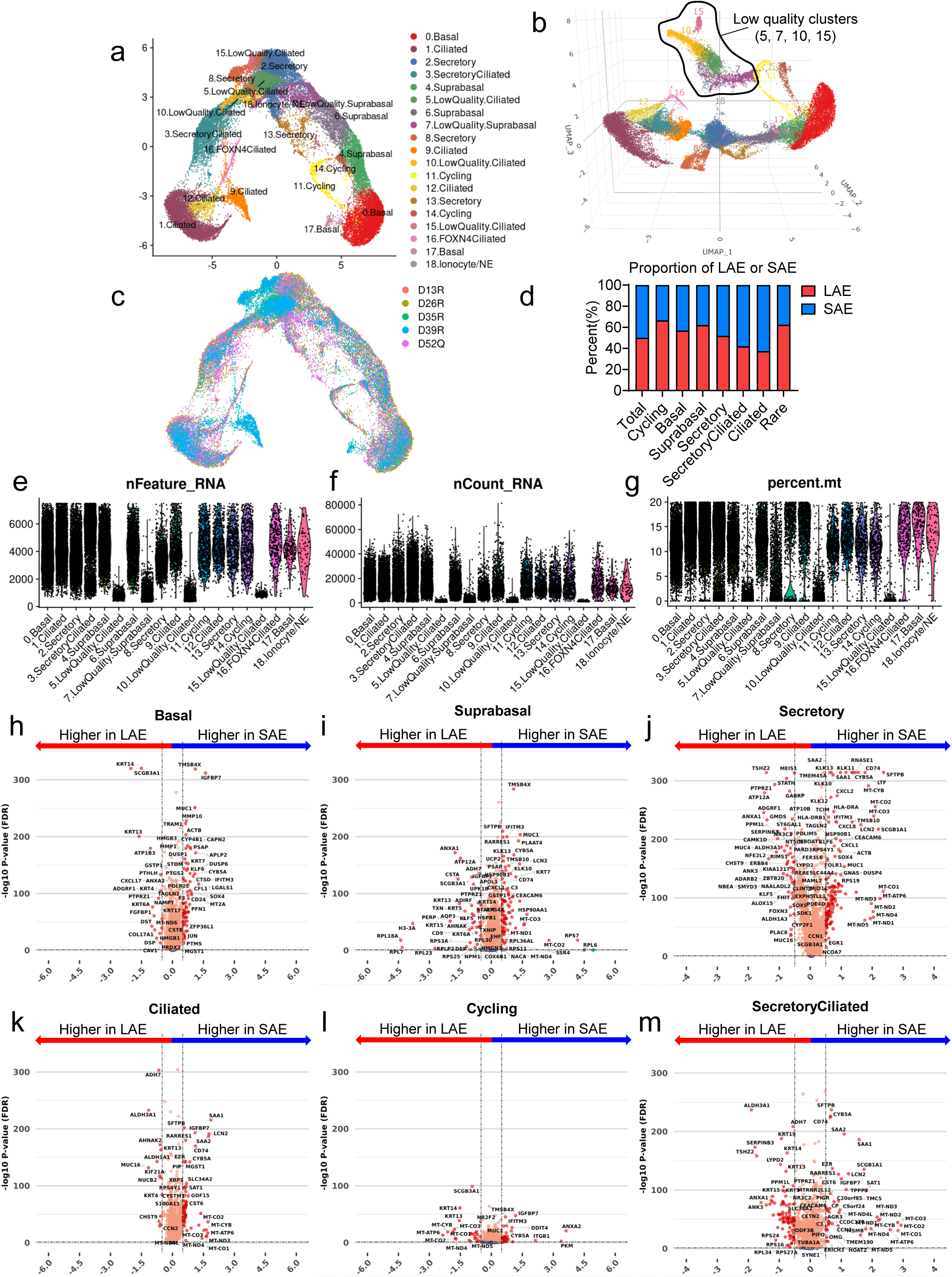
Single cell transcriptional profiling of LAE and SAE. **a, b.** UMAP of combined LAE and SAE cells showing 19 clusters in two-dimensional (**a**) and three-dimensional (**b**) space. **c.** UMAP colored by donor (n = 5). Clusters 5, 7, 10, and 15 were excluded based on low gene and transcript counts. **d.** Proportions of LAE and SAE cells across grouped clusters, normalized to total cell number per region. **e-g.** scRNA-seq quality metrics showing unique genes per cell (**e**), total transcript counts (**f**), and mitochondrial read percentage (**g**). Clusters 5, 7, 10, and 15 were considered as low-quality clusters and excluded from downstream analysis. **h-m.** Cell-type-specific differential gene expression between LAE and SAE cells within basal (**h**), suprabasal (**i**), secretory (**j**), ciliated (**k**), cycling (**l**), and secretory-ciliated (**m**) populations.

**Extended Data Fig. 4.**
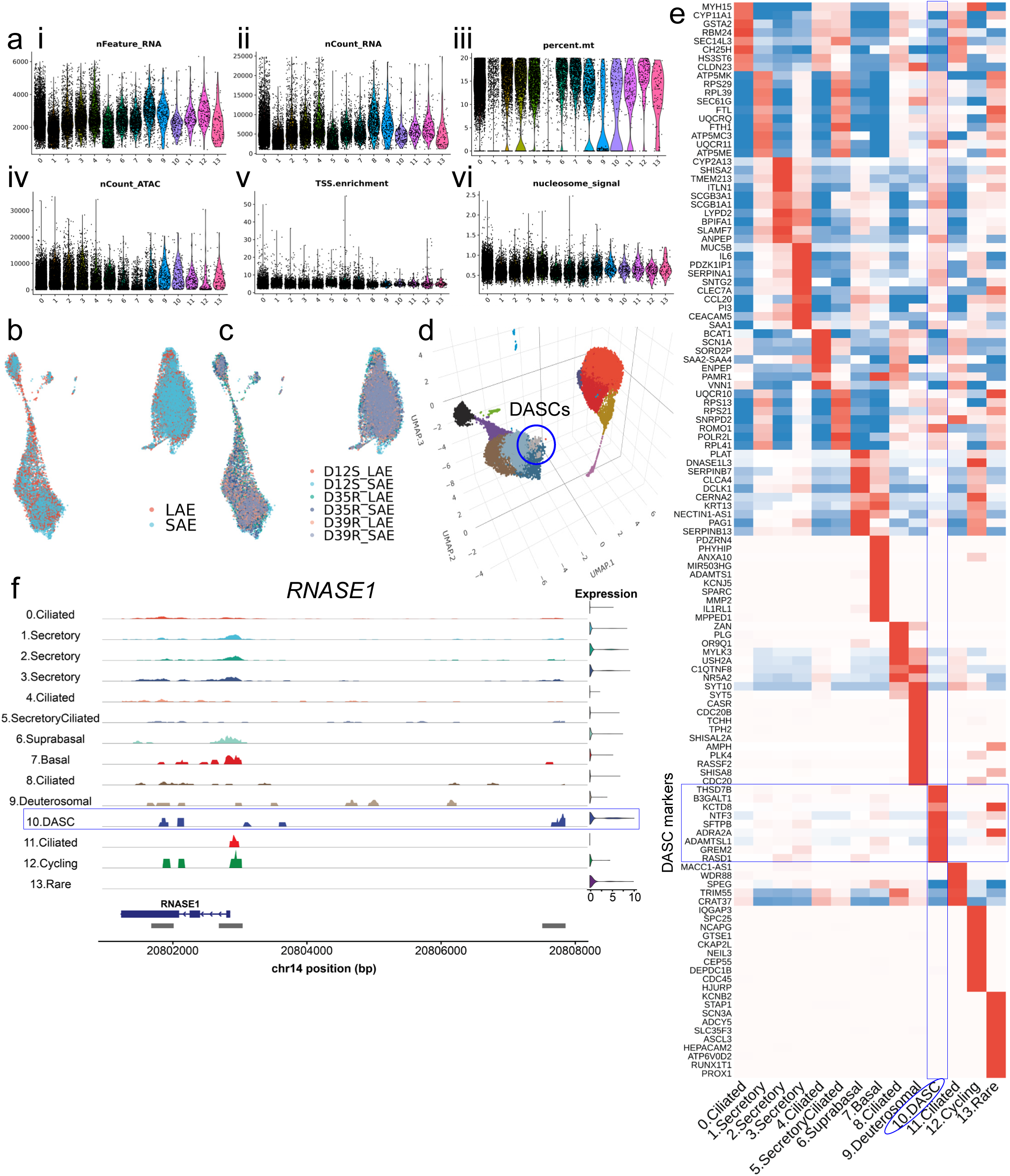
Integrated single-nucleus transcriptomic and chromatin accessibility profiling of LAE and SAE. **a.** snRNA/ATAC-seq quality metrics showing unique genes (**i**), transcript counts (**ii**), mitochondrial read percentage (**iii**), unique ATAC fragments (**iv**), transcription start site enrichment (**v**), and nucleosome signal (**vi**) per cluster. **b, c.** UMAP colored by region (LAE vs SAE) (**b**) and donor (n = 3) (**c**). **d.** Three-dimensional UMAP highlighting the DASC cluster. **e.** Heatmap of the top 10 marker transcripts per cluster. **f.** ATAC-seq peak tracks at the *RNASE1* locus showing DASC-enriched chromatin accessibility.

**Extended Data Fig. 5.**
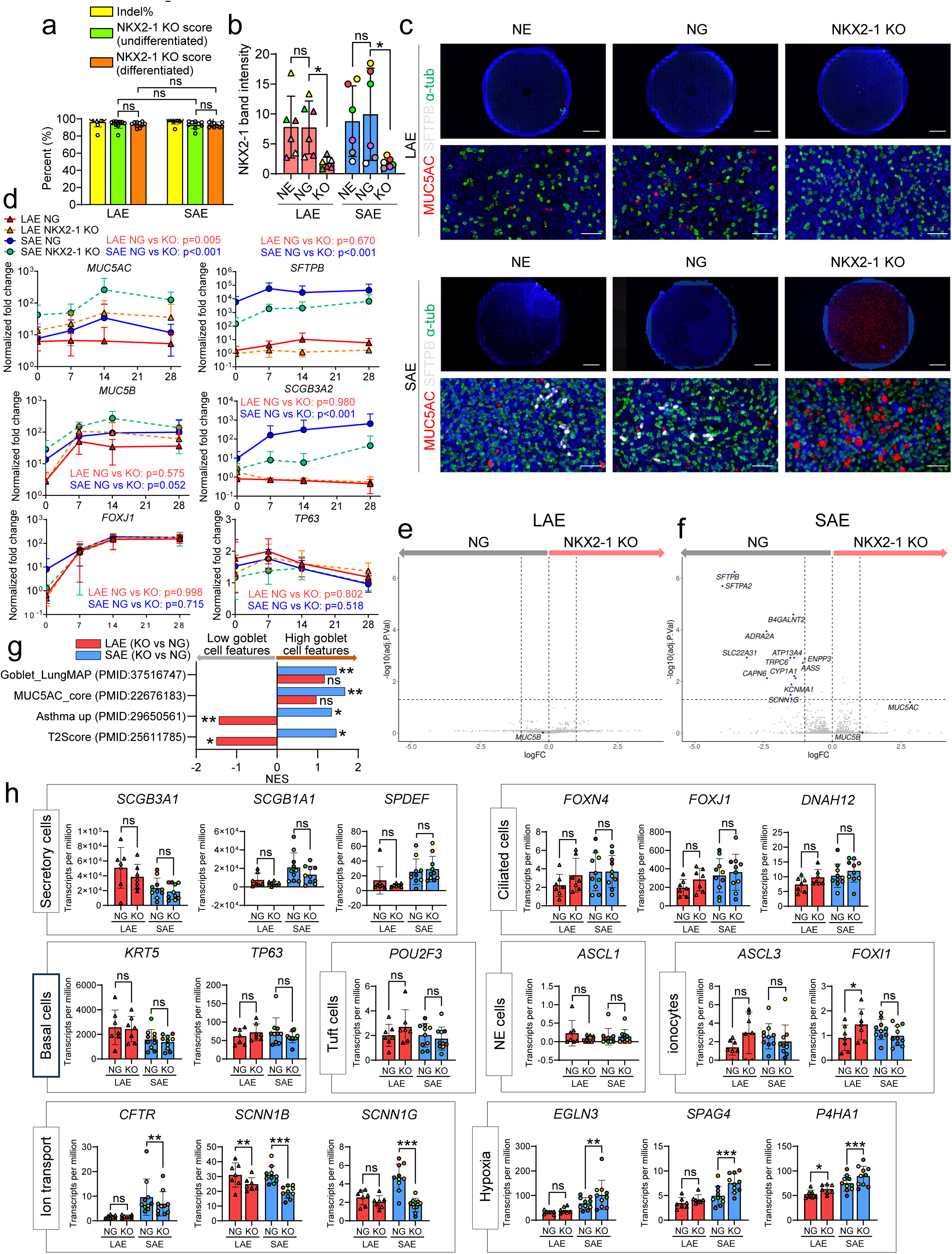
CRISPR/Cas9-mediated NKX2-1 knockout in LAE and SAE. **a.** Editing efficiency in LAE and SAE before and after differentiation, assessed by Inference of CRISPR Edits analysis (n = 10). **b.** Quantification of NKX2-1 protein by Western blot (n = 6). **c.** Immunofluorescence of non-electroporation control (NE), negative gRNA control (NG), or NKX2-1 KO LAE and SAE stained for MUC5AC, SFTPB, and α-tubulin. Scale bars, 2 mm (top) and 50 μm (bottom). **d.** Normalized fold changes in selected transcripts in NKX2-1 KO vs NG LAE or SAE during ALI differentiation. **e, f.** Differentially expressed genes in NKX2-1 KO vs NG LAE (**e**; n = 7) and SAE (**f**; n = 10), identified by bulk RNA-seq. **g.** Reactome pathway enrichment comparing NKX2-1 KO vs NG in LAE and SAE. **h.** Bulk RNA-seq expression of selected transcripts in NKX2-1 KO vs NG LAE and SAE; unadjusted p values are shown for predefined transcripts. ns, not significant; *p < 0.05; **p < 0.01; ***p < 0.001 (linear mixed-effects model).

**Extended Data Fig. 6.**
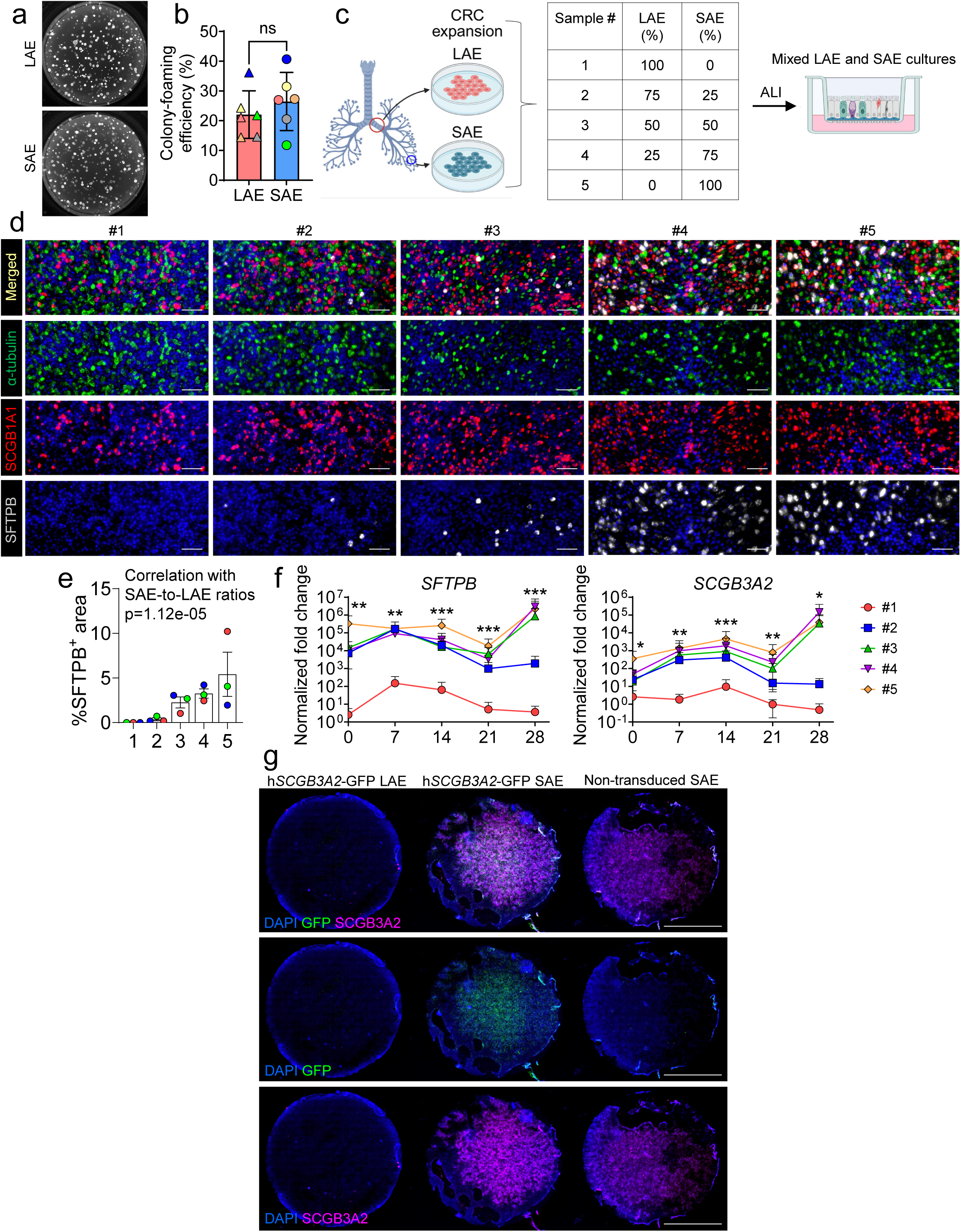
Differentiation properties of LAE and SAE progenitors. **a, b.** Representative colonies derived from LAE and SAE progenitors (**a**) and colony-forming efficiency in organoid cultures (n = 6) (**b**). **c.** Experimental design for mixed LAE and SAE progenitor cultures. **d, e.** Whole-mount immunofluorescence of fully differentiated mixed cultures (**d**) with quantification of SFTPB^+^ areas (**e**). Scale bar, 50 μm. Correlation between SFTPB^+^ area and SAE-to-LAE progenitor ratio was analyzed using a linear mixed model. **f.** qRT-PCR analysis of selected transcripts during ALI differentiation in mixed cultures; correlation with SAE-to-LAE progenitor ratio at each time point were assessed by a linear mixed model. **g.** Whole-mount immunofluorescence of h*SCGB3A2*-GFP-transduced LAE and SAE and non-lentiviral SAE controls after differentiation. Scale bar, 5 mm. ns, not significant; *p < 0.05; **p < 0.01; ***p < 0.001 (linear mixed-effect model).

**Extended Data Fig. 7.**
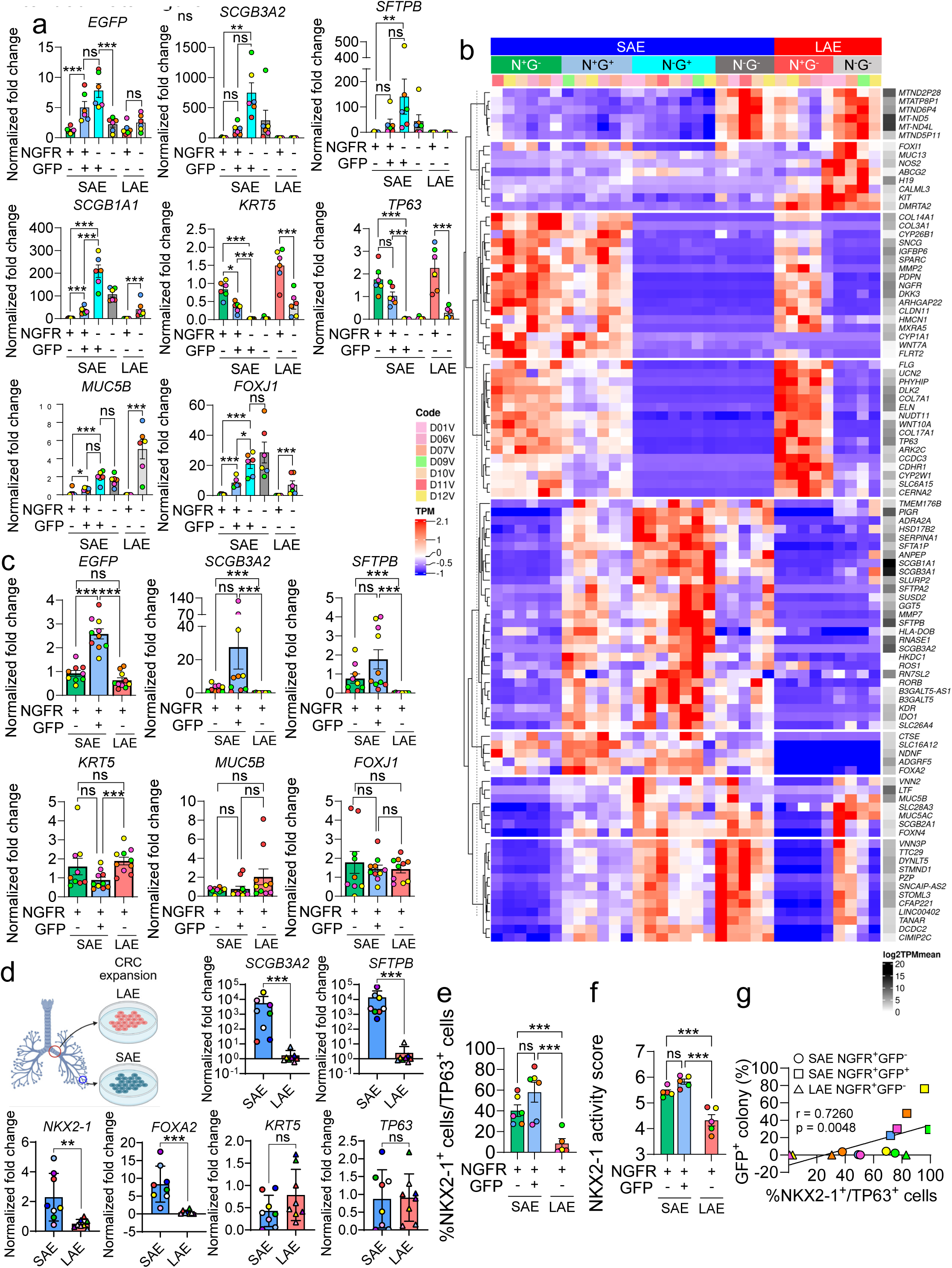
Transcriptional characterization of the distal airway epithelial lineage. **a.** qRT-PCR analysis of selected genes in sorted NGFR/GFP-defined populations from h*SCGB3A2*-GFP-transduced LAE and SAE. **b.** Heatmap of the top 20 differentially expressed genes across NGFR/GFP-defined populations. N, NGFR; G, GFP. **c.** qRT-PCR analysis of selected genes in 2D ALI cultures derived from FACS-sorted populations following CRC-expansion. **d.** qRT-PCR analysis of selected genes in CRC-expanded, undifferentiated LAE and SAE progenitors directly isolated from human lung tissue. **e.** Immunofluorescence-based quantification of NKX2-1^+^ cells among TP63^+^ basal cells in sorted NGFR^+^ basal subsets. **f.** NKX2-1 activity score inferred from regulatory network analysis of bulk RNA-seq data (n = 5). **g.** Spearman correlation between NKX2-1^+^ basal cell frequency among TP63^+^ basal cells and GFP^+^ organoid formation from corresponding sorted NGFR^+^ basal cell (BC) populations. ns, not significant; *p < 0.05; **p < 0.01; ***p < 0.001 (linear mixed-effect model).

**Extended Data Fig. 8.**
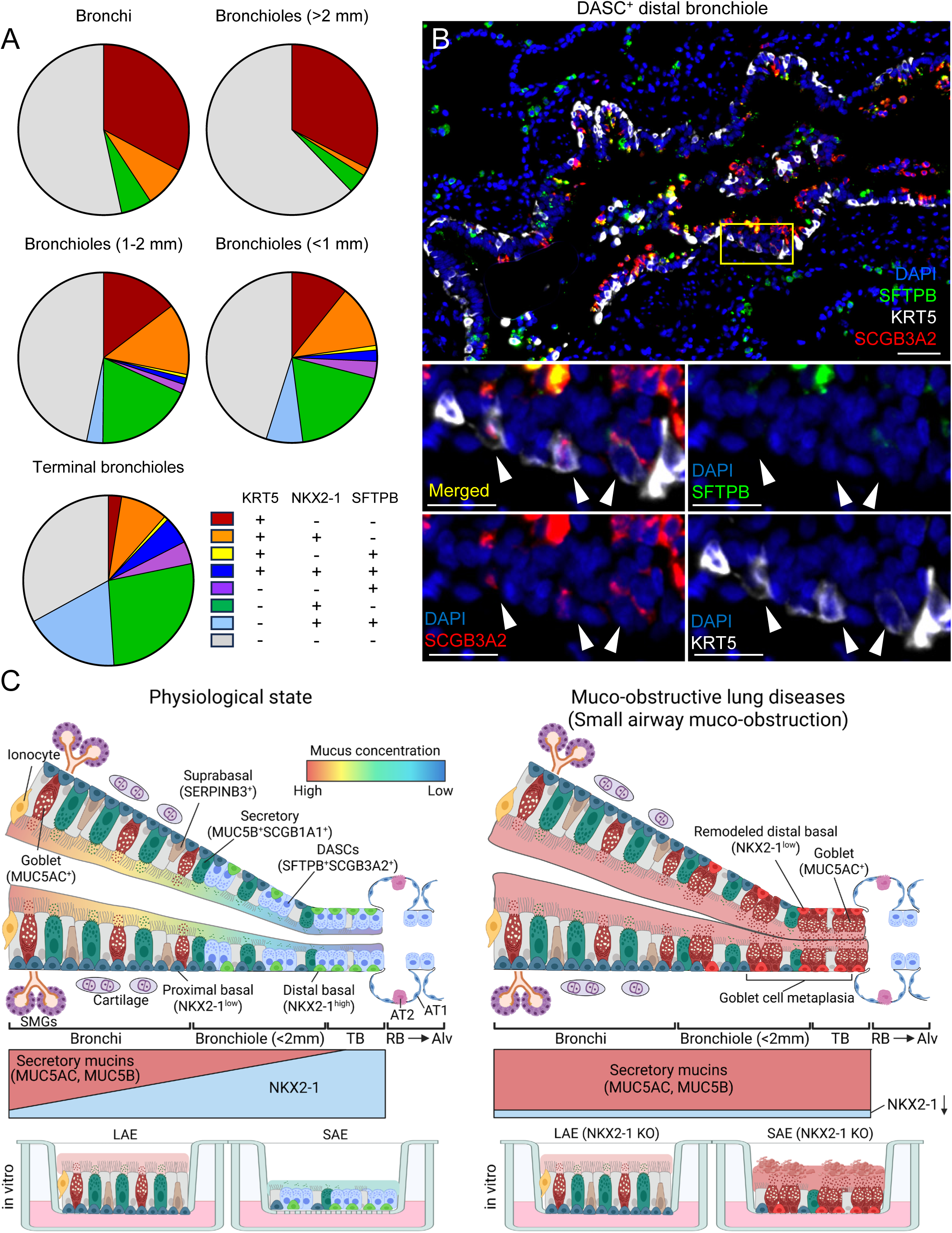
Localization of distal airway-specific basal cell subsets in human lungs. **a.** Quantification of epithelial subtypes defined by NKX2-1, KRT5, SFTPB across airway regions. TB, Terminal bronchiole. **b.** Immunofluorescence showing SCGB3A2^+^KRT5^+^ basal cells in a distal bronchiole of a normal human lung. Scale bar, 50 μm (20 μm insets). **c.** Schematic of NKX2-1-mediated regulation of mucus homeostasis in human distal airways in vivo and in vitro.

